# Inferring continuous and discrete population genetic structure across space

**DOI:** 10.1101/189688

**Authors:** Gideon S. Bradburd, Graham M. Coop, Peter L. Ralph

## Abstract

A classic problem in population genetics is the characterization of discrete population structure in the presence of continuous patterns of genetic differentiation. Especially when sampling is discontinuous, the use of clustering or assignment methods may incorrectly ascribe differentiation due to continuous processes (e.g., geographic isolation by distance) to discrete processes, such as geographic, ecological, or reproductive barriers between populations. This reflects a shortcoming of current methods for inferring and visualizing population structure when applied to genetic data deriving from geographically distributed populations. Here, we present a statistical framework for the simultaneous inference of continuous and discrete patterns of population structure. The method estimates ancestry proportions for each sample from a set of two-dimensional population layers, and, within each layer, estimates a rate at which relatedness decays with distance. This thereby explicitly addresses the “clines versus clusters” problem in modeling population genetic variation. The method produces useful descriptions of structure in genetic relatedness in situations where separated, geographically distributed populations interact, as after a range expansion or secondary contact. We demonstrate the utility of this approach using simulations and by applying it to empirical datasets of poplars and black bears in North America.

**Author summary:** One of the first steps in the analysis of genetic data, and a principal mission of biology, is to describe and categorize natural variation. A continuous pattern of differentiation (isolation by distance), where individuals found closer together in space are, on average, more genetically similar than individuals sampled farther apart, can confound attempts to categorize natural variation into groups. This is because current statistical methods for assigning individuals to discrete clusters cannot accommodate spatial patterns, and so are forced to use clusters to describe what is in fact continuous variation. As isolation by distance is common in nature, this is a substantial shortcoming of existing methods. In this study, we introduce a new statistical method for categorizing natural genetic variation - one that describes variation as a combination of continuous and discrete patterns. We demonstrate that this method works well and can capture patterns in population genomic data without resorting to splitting populations where they can be described by continuous patterns of variation.

## Introduction

A fundamental quandary in the description of biological diversity is the fact that diversity shows both discrete and continuous patterns. For example, reasonable people can disagree about whether two populations are separate species because the process of speciation is usually gradual, and so there is no set point in the continuous divergence of populations when they unambiguously become discrete species. The issue of identifying meaningful biological subunits extends below the species level, as patterns of phenotypic and genetic diversity within and among populations are shaped by continuous migration and drift, as well as by more discrete events, such as geographic barriers, rapid expansions, bottlenecks, and rare long-distance migration. Both discrete and continuous components are required to accurately describe most species’ patterns of genetic relatedness.

From a practical standpoint, even while acknowledging continuous processes, we often need to identify somewhat separable populations from which individuals are sampled [1]. Delineating populations is useful for systematics and for informing conservation priorities [2–4]. Furthermore, we often need to identify subsets of indivduals resulting from reasonably coherent evolutionary histories for downstream analyses to learn about population history and adaptation. Conversely, the substantial information available from continuous, geographic differentiation (e.g., adaptation along a climatic gradient) can be confounded by discrete historical processes (e.g., admixture), requiring methods that can disentangle the two.

There have been many methods proposed to characterize population genetic structure, including generating population phylogenies [5, 6], dimensionality-reduction approaches, such as principal components analysis [7–10], and model-based clustering approaches (e.g., [11–20]), among others. Each of these methods perform best in particular situations, but many can give misleading results when applied to data that show a continuous pattern of differentiation, as that produced by geographic isolation by distance [9, 21, 22]. Here, we will focus on model-based clustering, the most widely used class of approaches for population delineation. Existing model-based clustering methods model each individual’s genotypes as random draws from a set of underlying, unobserved population clusters, each with a characteristic set of allele frequencies, which are estimated. These underlying frequencies are identical for all individuals assigned to a cluster, *regardless* of their spatial location. Spatial information has been incorporated into some of these methods, by, for example, placing spatial priors on cluster memberships [19,20], but this does not address the underlying issue that these methods assume that allele frequencies are constant in a cluster across the species’ range.

*Isolation by distance* (IBD) refers to a pattern of increasing genetic differentiation with geographic separation, which occurs when geographically restricted dispersal allows genetic drift to build up differentiation between distant locations [21]. Theoretical work, mostly derived from “stepping-stone” models [23–25], gives us some analytical predictions for isolation by distance [26–28], but substantial work remains to be done [29,30]. Given the generality of the circumstances that generate a pattern of isolation by distance, it is unsurprising that isolation by distance is very widespread in nature [31,32].

The ubiquity of isolation by distance presents a challenge for models of discrete population structure, as it is frequently difficult to determine whether observed patterns of genetic variation are continuously distributed across a landscape, or instead are partitioned in discrete clusters. This problem can be compounded if sampling is done unevenly or discretely across a population or species’ range, and has given rise to a debate in the population genetic literature about how best to describe sets of individuals using continuous clines and discrete clusters (e.g., [33, 34]).

Existing model-based clustering approaches can only describe continuous patterns of variation using discrete clusters, and so tend to erroneously describe continuously distributed variation with multiple clusters that show spatially autocorrelated cluster membership [22, 31]. In analyses of empirical datasets, which often show strong isolation by distance, model-based clustering approaches will therefore tend to overestimate the number of discrete clusters present.

To address this, we set out to develop a model-based clustering method that, when possible, uses isolation by distance to explain observed genetic variation. With an explicit spatial component, discrete population structure need only be invoked when genetic differentiation in the data deviates significantly from that expected given geographic separation. In this paper, we model genetic variation in genotyped individuals as partitioned within or admixed across a specified number of discrete layers, within each of which relatedness decays as a parametric function of the distance between samples. We also implement a cross-validation approach for comparing and selecting models across different numbers of layers, and we demonstrate the utility of our approach using both simulated and empirical data. The implementation of this method, `conStruct` (for “*continuous structure*”), is available as an R package at https://github.com/gbradburd/conStruct.

## Results

### Data

The statistical framework of our approach is conceptually similar to that in [35], [36], and [37], although we use a somewhat different summary statistic than in this previous work. The model works with allele frequencies at *L* unlinked, bi-allelic single nucleotide polymorphisms (SNPs) genotyped across *N* samples. Each “sample” may be a single individual, a collection of individuals from a location, or frequencies estimated from pooled sequencing. We write *N*_*i*_ for the number of chromosomes sequenced in the *i*^*th*^ sample. The *sample frequency* at locus *l* in sample *n*, denoted *f*_*n,l*_, is calculated by first arbitrarily choosing one of the observed alleles at locus *l* to count, then dividing the number of observations of that counted allele by the total number of chromosomes genotyped at that locus in sample *n*. The choice of allele does not affect subsequent calculations, and so may be arbitrary. We then calculate the *allelic covariance* between samples *i* and *j*, denoted 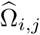, as the expected covariance of distinct individual alleles chosen from each of the two samples at a random locus, coding the counted allele as ‘1’ (and the other as ‘0’). Concretely, this is

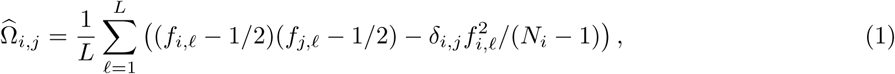

where *δ*_*i,j*_ = 1 if *i* = *j* and is 0 otherwise. Although we describe this as a covariance between individually drawn alleles, 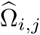 is in fact also the covariance between the allele frequencies of a randomly chosen allele in samples *i* and *j*, as long as *i* ≠ *j*. The diagonal (where *i* = *j*) does not have this interpretation, and reflects the covariance between individual alleles drawn from within the population.

This definition of covariance differs from the usual “genetic covariance” [38] in that (a) we do not subtract locus means, and so to make the statistic invariant under choice of reference allele, (b) we randomly choose an allele to count at each locus. We discuss the derivation of Eq. (1) further in “Allelic covariance”. As noted in [39], this covariance has a close relationship to the pairwise genetic distance: if π_*i,j*_ is the mean density of sites at which random samples from *i* and *j* differ at a randomly chosen locus, then Ω_*i,j*_ = (1 – 2π_*i,j*_)/4. Therefore, this allelic covariance is more affected in shape by singleton sites than the standard genetic covariance, so it may be advisable to filter these prior to analysis if they are likely to contain a large percentage of errors [40].

### Continuous and discrete differentiation

Clustering approaches to describing genetic variation are useful because population history can often be meaningfully described on a coarse scale by interactions between discrete “populations” whose relationships are delimited by patterns of glaciation, large-scale migration, mountain ranges, and the like. Here we add a spatial component within each such discrete historical component, which we refer to as a set of “layers” that overlay the modern map. We imagine each layer as a geographically distributed population that extends over the entire sampled range of the populations. As depicted in Figure 1, each sample is composed of a mixture of contributions from each of these layers, with the relative contributions of each layer described by a set of “admixture proportions” (the 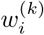). These layers thus take the place of “clusters” in clustering methods, but we do not adopt this term, as “spatial cluster” suggests a clustering in space, while our layers may contribute to genetic variation across the entire geographic range.

**Fig 1.**
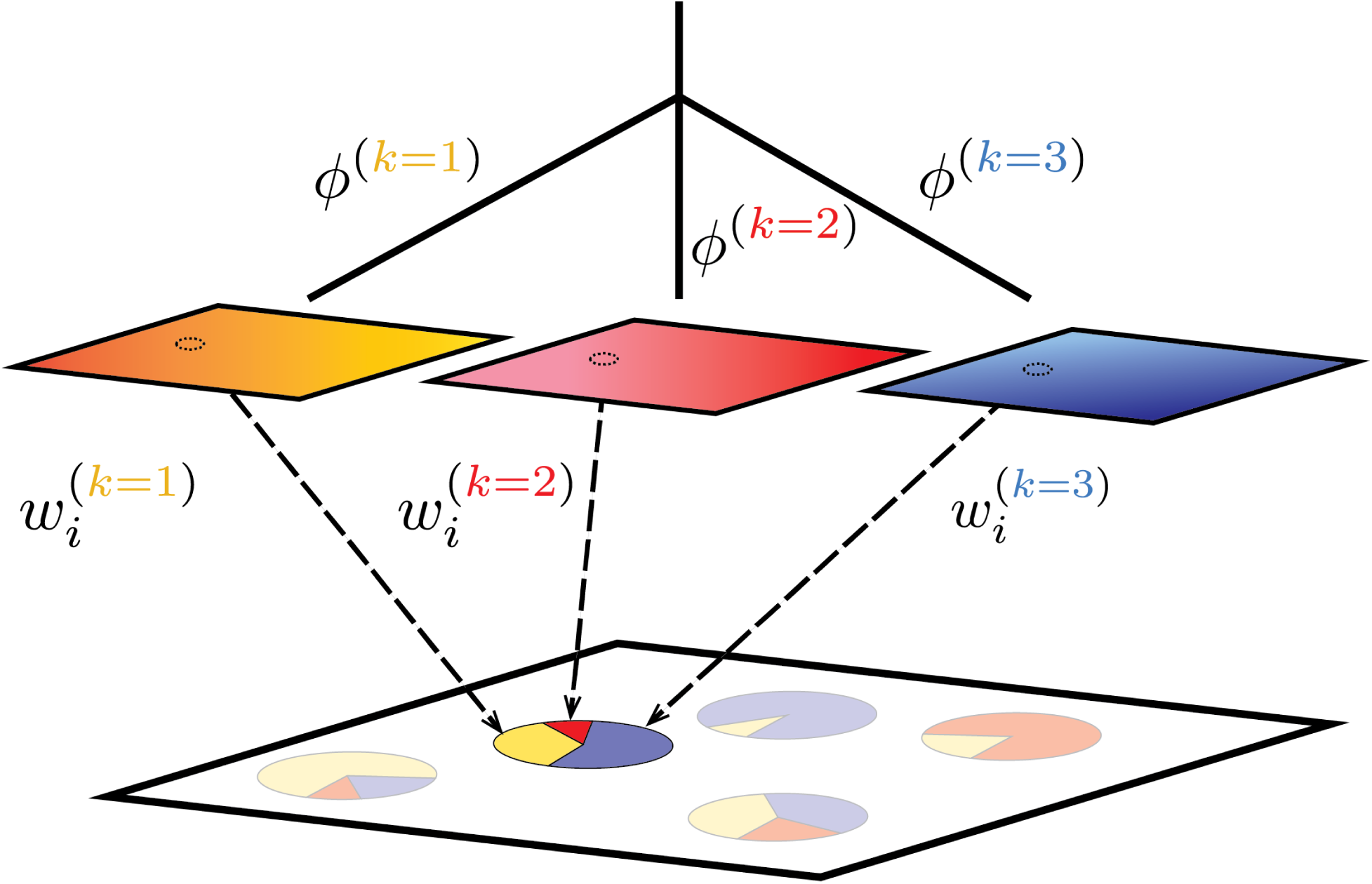
Schematic of our method, using *K* = 3 as an example. Spatial autocorrelation of allele frequencies within each layer is depicted by color gradients, and *ϕ*^(*k*)^ denotes the covariance shared by samples with ancestry entirely in the *k*th layer. Sampled populations on the landscape are inferred to be admixed between these layers; the *i*th sample draws proportion 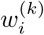 of its ancestry from layer *k*. For convenience, each layer is depicted as a small square, but in fact, each layer exists everywhere in the sampled area, so the small dashed circles on each layer show where the location of the highlighted admixed sample intersects each layer.

Within each of these layers, allele frequencies have positive covariance at geographically close locations, but this covariance is allowed to decay as geographic distance increases. This pattern of spatial decay reflects how migration between neighboring denies homogenizes allele frequency changes that arise locally due to drift, but less effectively homogenizes geographic distant demes, resulting in a continuous pattern of isolation by distance within each layer. There is a fixed amount of covariance between layers, irrespective of spatial location. Within each layer, allele frequencies are expected to change gradually with distance, but observed frequencies can change abruptly at many loci if the proportions of ancestry individuals derive from each layer (the “admixture” proportions) do so as well.

To allow flexibility in the form of the decay of allelic covariance with geographic distance within each layer, we define the covariance within layer *k* between samples *i* and *j* to be:

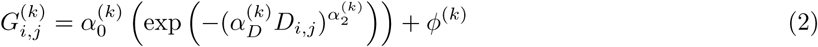

where the superscript (*k*) denotes parameters specific to the *k*th layer. The quantity *D*_*i,j*_ is the observed geographic distance between samples *i* and *j*, and the *α*^(*k*)^ parameters control the shape of the decay of covariance with distance in the layer. Our choice of a powered-exponential decay, as parameterized by the αs, is a flexible and standard choice in spatial statistics [41], and is not chosen to match a particular population genetics model. The *ϕ*^(*k*)^ is a parameter that describes the background covariance within the layer. If two samples draw 100% of their ancestry from layer *k*, then their covariance under the model is 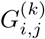; if they are furthermore geographically very close (*D*_*i,j*_ = 0) they will have covariance 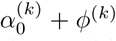 If the geographic distance between them is very large, their covariance will be equal to the background level *ϕ*^(*k*)^ within the layer. The “shared drift” parameter *ϕ*^(*k*)^ is analogous to the branch length connecting the *kth* population to the population ancestral to all modeled layers (see for example [42,43]), although they cannot be directly compared because we are modeling the allelic, rather than genetic, covariance. In “Model rationale” we lay out a simple model of allele frequencies underlying this covariance model.

We then allow samples to draw their ancestry from more than one layer. The “admixture” proportion of the *i*th sample in the *k*th layer, denoted 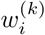, gives the genome-wide proportion of alleles from sample *i* that derive from layer *k* (and so 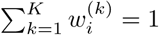). A visual representation of the method is shown in Fig 1.

We can then describe the covariance between samples *i* and *j* across all *K* layers, Ω_*i,j*_, by summing their within-layer spatial covariances (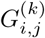 in layer *k*), weighted by the relevant admixture proportions.

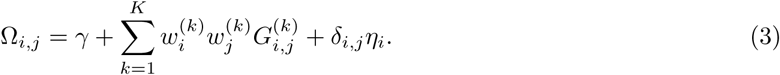

In this equation, 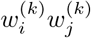 is the proportion of alleles that *both* sample *i* and sample *j* have inherited from layer *k*.

In addition to the admixture-weighted sum of the within-layer spatial covariances, this function contains two terms, γ and *δ*_*i,j*_*η*_*i*_. The first, γ, describes the global allelic covariance between all samples, and arises because all samples share an ancestral mean allele frequency at each locus, which generates a base-line covariance. In the final term, *δ*_*i,j*_ is an indicator variable that takes a value of 1 when *i* equals *j* and 0 otherwise, and *η*_*i*_ adds variance specific to sample *i*. This term on the diagonal of the parametric covariance matrix captures processes shaping variance within the sampled deme, such as inbreeding and the sampling process.

### Likelihood and inference

If the allele frequency deviations at each locus were independent between loci and multivariate normally distributed across populations, their allelic covariance 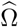 would be Wishart distributed with degrees of freedom equal to *L*, the number of loci genotyped. We use this as a convenient approximation to the true distribution described above, and so define the likelihood of the allelic covariance to be

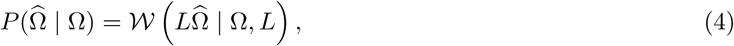

where 𝒲 is the Wishart likelihood function. Statistical nonindependence between loci (linkage disequilibrium) will decrease the effective number of degrees of freedom. One possible solution, which we have not yet found necessary to implement, would be to estimate an *effective* number of loci by introducing a parameter to modify the given degrees of freedom and thereby informally model linkage between loci (e.g., [39]).

We estimate the values of the parameters of the model using a Bayesian approach. Acknowledging the dependence of the parametric covariance matrix Ω on its constituent parameters *w*, *α*, *ϕ*, *η*, γ and on the (observed) geographic distances *D* with the notation Ω(*w*, *α*, *ϕ*, *η*, γ, *D*), we denote the posterior probability density of the parameters as:

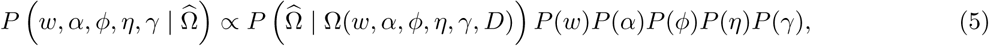

where *P*(*w*), *P*(*α*), *P*(*ϕ*), *P*(*η*), and *P*(γ), are prior distributions. All parameters are given (half-)Gaussian priors except for *α*_2_ which is uniform on (0, 2), and *w*, for which we use an independent Dirichlet of dimension *K* for each sample (see Table 1 for specifics). Parameters are independent between layers. We use Hamiltonian Monte Carlo as implemented in STAN [44–47] to estimate the posterior distribution on the parameters. Our R package, `conStruct` (for “*continuous structure*”), functions as a wrapper around this inference machinery.

**Table 1.** List of parameters used in the `conStruct` model, along with their descriptions and priors. The mean of the Normal prior on γ, 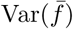, is the variance of the sample mean allele frequencies across loci.

| Parameter | Description | Prior |
| --- | --- | --- |
| $\gamma$ | global covariance due to shared ancestral frequency | $\gamma \sim \mathcal{N}(\mu = \text{Var}(\bar{f}), \sigma = 0.5)$ |
| $\alpha_0^{(k)}$ | controls the sill of the covariance matrix in layer $k$ | $\alpha_0^{(k)} \sim \mathcal{N}(\mu = 0, \sigma = 1)$ |
| $\alpha_D^{(k)}$ | controls the rate of the decay of covariance with distance in layer $k$ | $\alpha_D^{(k)} \sim \mathcal{N}(\mu = 0, \sigma = 1)$ |
| $\alpha_2^{(k)}$ | controls the shape of the decay of covariance with distance in layer $k$ | $\alpha_2^{(k)} \sim U(0, 2)$ |
| $\eta_i$ | the nugget in population $i$ (population specific drift parameter) | $\eta_i \sim \mathcal{N}(\mu = 0, \sigma = 1)$ |
| $\phi^{(k)}$ | layer-specific shared drift in layer $k$ | $\phi^{(k)} \sim \mathcal{N}(\mu = 0, \sigma = 1)$ |
| $w_i$ | admixture proportions sample $i$ draws across $K$ layers | $w_i \sim \text{Dir}(\alpha_1 \dots \alpha_K = 0.1)$ |

### Relationship of this model to nonspatial structure models

A nice feature of our approach is that the model described in Eq. (3) contains a nonspatial assignment model as a special case (see “Models, parameters, and priors” for a more in-depth discussion). By setting 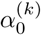 to zero for all *k*, we obtain a nonspatial model in which each cluster has its own allele frequency at each SNP, and individuals draw a proportion of their ancestry from each cluster. This model is very similar to that of STRUCTURE [11] and related models (e.g., [14]); the main difference is that our likelihood assumes that allele frequencies are normally distributed around their expectations, while the standard assignment methods assume that the error is binomially distributed [48]. (We make this approximation for the substantial advantages in computational speed.) The second small difference is that, in the original STRUCTURE model, allele frequencies at each locus are independently drawn for each cluster [11], while in conStruct’s non-spatial model, it is more natural to envision each cluster’s allele frequency as being drifted away from a single, global allele frequency. This makes our model more closely related to the “*F*-model” prior for allele frequencies of [12]. As these differences are relatively small, we can compare the fit of the different models — spatial vs. nonspatial, across different values of *K* — by comparing their performance in a common framework. We also validate our claim that the results of our nonspatial method are a close match to those of STRUCTURE-based approaches.

### Choice of layer number and cross-validation

There are a number of reasons why there is no true (or right) number of layers for real datasets, discussed further in the Discussion. However, it is still important to assess whether additional layers (larger *K*) meaningfully model patterns in the data or merely explain spurious variation introduced by noise - in other words, whether additional model complexity provides significant explanatory power. Toward that end, we have implemented a method for statistically comparing `conStruct` results across different values of *K* and between the spatial and nonspatial models.

Several approaches have been used as model choice criteria for the number of discrete clusters in population genetic data, including: comparisons of the likelihood of the data across different values of *K*, with various criteria on how to choose a single value (e.g., [49]), or with information theoretic penalizations such as AIC or BIC (e.g., [14]); or comparisons of the marginal likelihood, generated either via various approximations (e.g., [11]) or via a more rigorous and computationally intensive approach such as thermodynamic integration [50] or inference using a Dirichlet process prior [17]. See [50] for a discussion of these approaches and comparison between several methods.

We use cross-validation (similar in spirit to [51]) to attack this problem. To do this, we use a “training” partition of the data (in practice, a random 90% subset of the loci) to estimate the posterior distribution of the parameters, and then calculate the log-likelihood of the remaining “testing” loci, averaged over the posterior. Prediction accuracy of a particular model (e.g., value of *K*) is then measured using this log-likelihood, averaged over a number of independent data partitions. The best model is judged to be the simplest one with significantly better predictive accuracy than others (see “Cross validation” for more on our cross-validation procedure). In general, larger values of *K* allow the model more flexibility, and thus increases the likelihood of the training partition, but this improvement in the likelihood will plateau (or even peak), as above a certain *K* the model only fits noise specific to the training data rather than generalizable patterns. At any value of *K*, support for the spatial model over the nonspatial model means that isolation by distance is likely a feature of the data.

Cross-validation provides a valuable summary of how much explanatory power is added by spatial structure within each layer, and each additional layer. However, we remind users that “statistical significance does not imply real-world significance’,’ and so small but statistically significant differences between models should probably not be relied on too strongly.

Another way to describe the practical significance of additional layers is to calculate each layer’s relative contribution to total covariance, and to choose a value of *K* where all layers have a contribution above some cutoff (e.g. 0.1%). The Dirichlet prior on admixture proportions is quite harsh against intermediate admixture values (see Table 1), encouraging the model to “not use” unnecessary layers if they are present in the model, so that they will have a low contribution to overall covariance.

To calculate layer contributions, we use the following alternative description of our covariance model: the genomes of any pair of individuals *agree* with some background probability at a locus, but this probability of agreement is increased on any segment of genome that both have inherited from the same layer (the amount it increases depends on how far apart they are geographically and on the decay of isolation by distance). We use this characterization to quantify the relative contributions of each layer, by computing the average contribution to increased probability of agreement as described in “Calculating layer contributions”). This layer contribution is similar to the “ancestry contribution” proposed by [16]. However, each of our layers can induce a different amount of covariance between samples embedded in them, so we take that into account when calculating each layer’s contribution to the whole.

## Simulations

To test the method, we first generated data using the coalescent simulator ms [52]. In each simulation, we split a single ancestral population into *K* subpopulations *τ*_s_ units of coalescent time in the past, and at time *τ*_e_ in the past, each of these discrete populations instantaneously colonized a separate 6 × 6 square lattice of demes. Migration on each lattice was to nearest neighbors (eight neighbors, including diagonals). Finally, at time *τ*_a_ in the past, we collapsed those *K* discrete layers into a single grid of demes, choosing various amounts of admixture from these different layers (see Fig 10). We collapsed the layers together using random, spatially autocorrelated admixture proportions (see “Simulation details”). We simulated datasets using *K* = 1, 2, and 3 layers; in each simulation we sampled 10,000 unlinked loci from each of 20 haploid individuals from every deme. We then ran both spatial and nonspatial `conStruct` analyses on each simulated dataset with *K* between 1 and 7, and compared predictive performance of the models using cross-validation. For comparison, we also analyzed each simulated dataset using fastSTRUCTURE [16], using *K* = 2 – 4.

**Fig 10.**
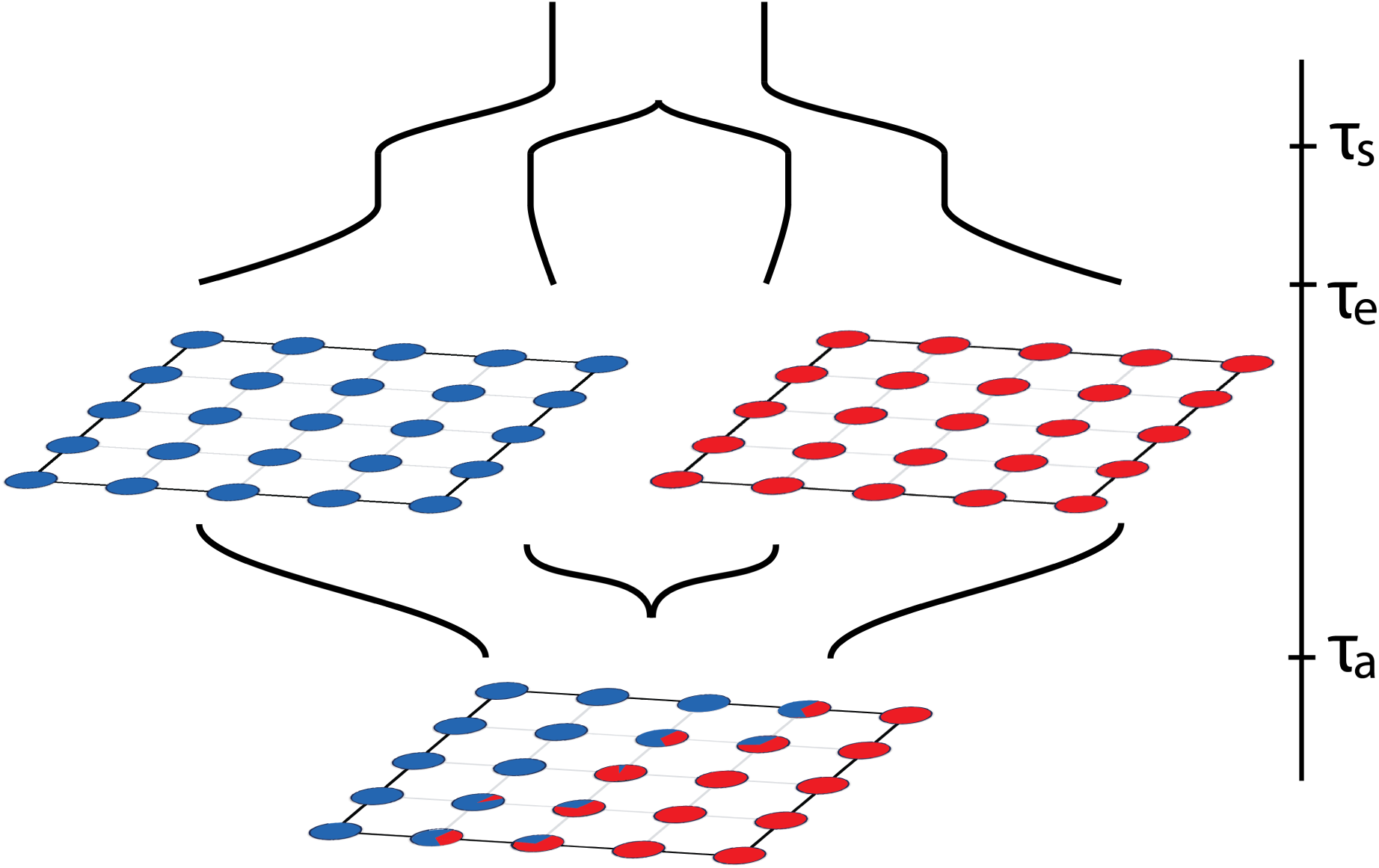
Schematic of how we simulate datasets with continuous and discrete differentiation, using *K* = 2 as an example. Going forward in time, the *K* populations split from a common ancestor at time *τ*_s_, then expand to each colonize a lattice of demes with nearest-neighbor symmetric migration at time *τ*_e_, then finally at time *τ*_a_ collapse into a single lattice consisting of demes with ancestry entirely in one or the other of the populations, or admixed between them.

With these simulations, spatial `conStruct` does not create spurious discrete groupings when there are none: Figures 2, S5, and S10 show that subsequent layers beyond the number used for simulation are unused. When data are simulated with *K* = 1 but analyzed with *K* > 1, the layers are present, but contribute very little to any population. Even when the spatial model is run with *K* = 7, the inferred admixture proportions are nearly identical to those estimated under the true value of *K* for each simulation. Moreover, the method infers the true admixture proportions with high accuracy, tight precision, and good coverage (Figs S8 and S13).

In contrast, the nonspatial model describes geographic variation using gradients of admixture between more and more discrete clusters to better approximate the continuous, spatial patterns of relatedness (depicted in Figs 2 and S1). The fastSTRUCTURE results are qualitatively similar, as shown in Figs S14–S16. Each nonspatial cluster is in truth genetically more similar within itself than it is to other clusters, but we know that these boundaries are arbitrary, because the data were simulated without them.

**Fig 2.**
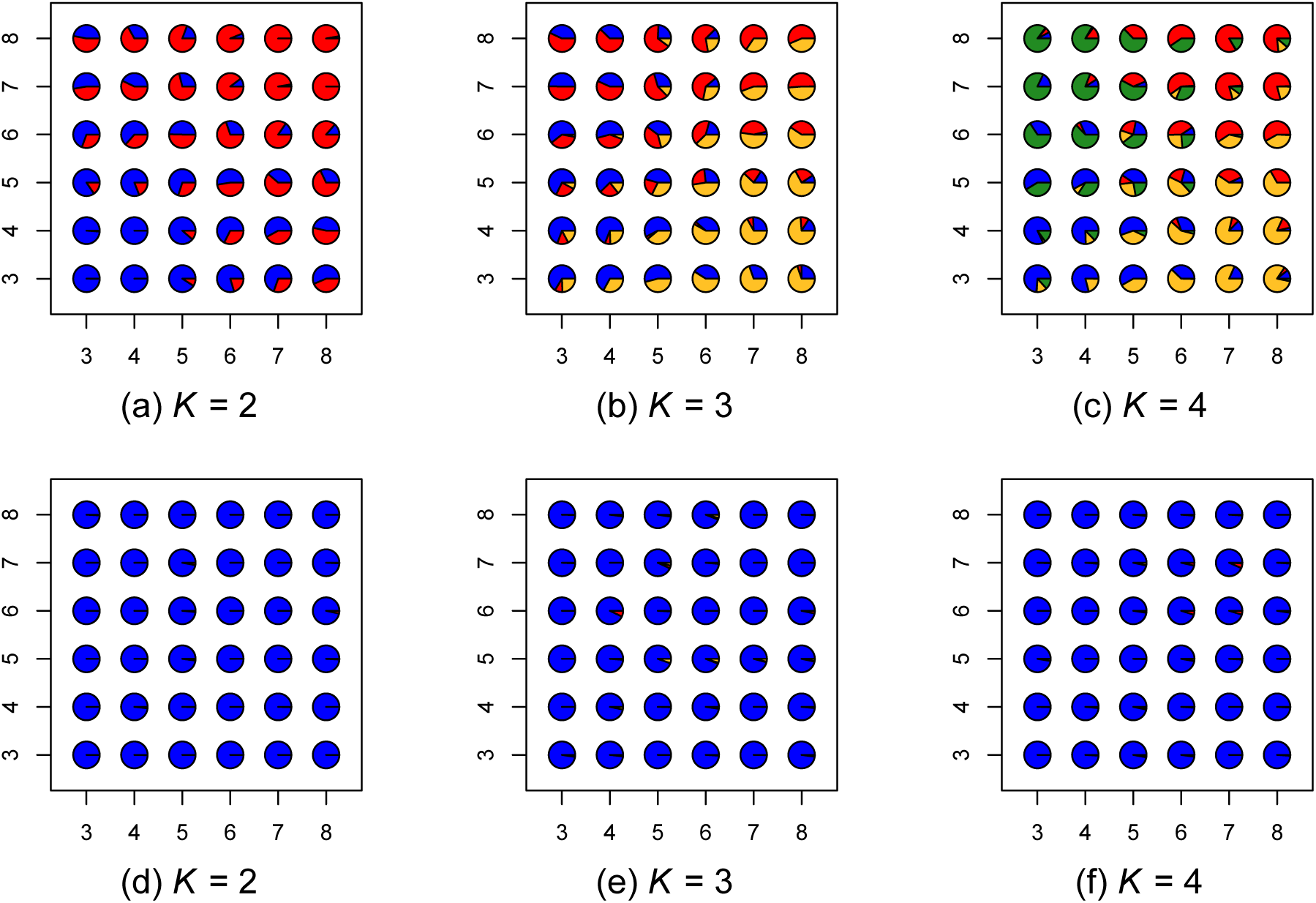
Results for data simulated using *K* = 1, showing maps of admixture proportions estimated using the nonspatial model for *K* = 2 through 4 (top row) and the spatial model for *K* = 2 through 4 (bottom row). Since there is only a single layer in the simulation, the spatial model accurately depicts the data in all cases, while the nonspatial model creates spurious clusters.

The spatial model’s better fit is reflected by increased predictive accuracy: as shown in Fig 3, across all models and choices of *K*, the spatial model is correctly preferred over the nonspatial model. As desired, predictive accuracy of the spatial model increases until the true value of *K*, and then plateaus or declines (Figs 3, S3, S7, and S12), while predictive accuracy of the nonspatial model increases as subsequent clusters are added.

**Fig 3.**
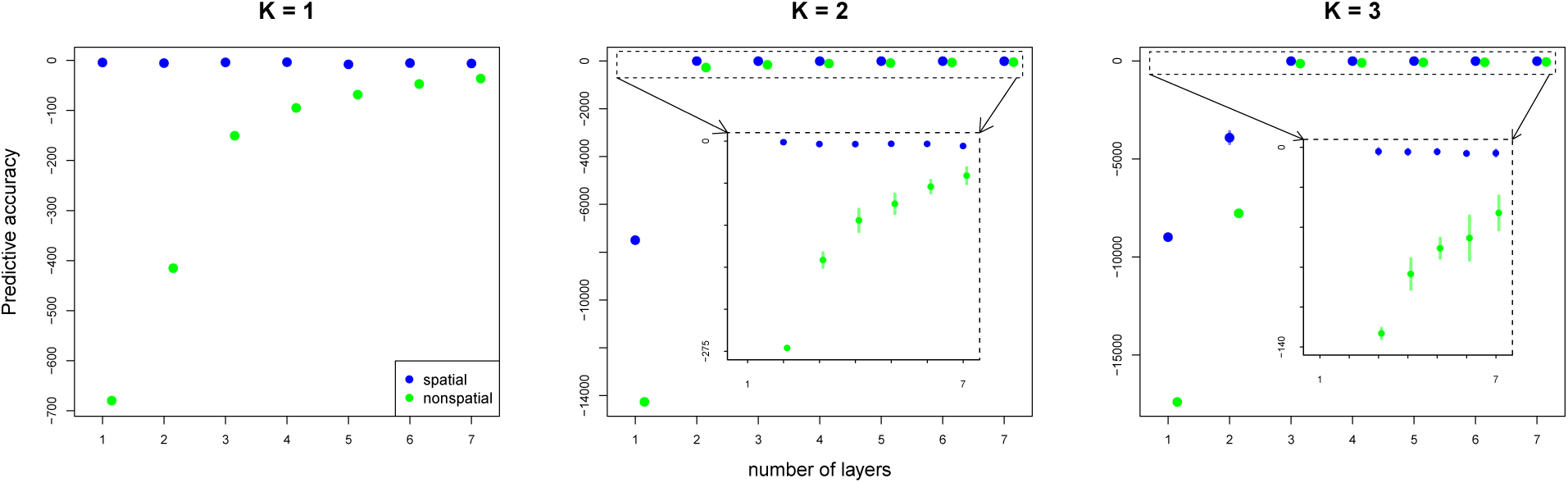
Cross-validation results for data simulated under *K* = 1, *K* = 2, and *K* = 3, comparing the spatial and nonspatial `conStruct` models (in blue and green, respectively) run with *K* = 1 through 7. The inset plots zoom in on cross-validation results outlined in the dotted boxes. The spatial model shows better model fit at every value of *K*.

The unimportance of spurious layers can be seen in plots of layer contributions (Figs 4, S6, and S11). In the spatial analyses, once we pass the true *K*, subsequent layers add little in terms of (co)variance explained; in contrast, additional clusters in the nonspatial analyses continue to contribute substantially.

**Fig 4.**
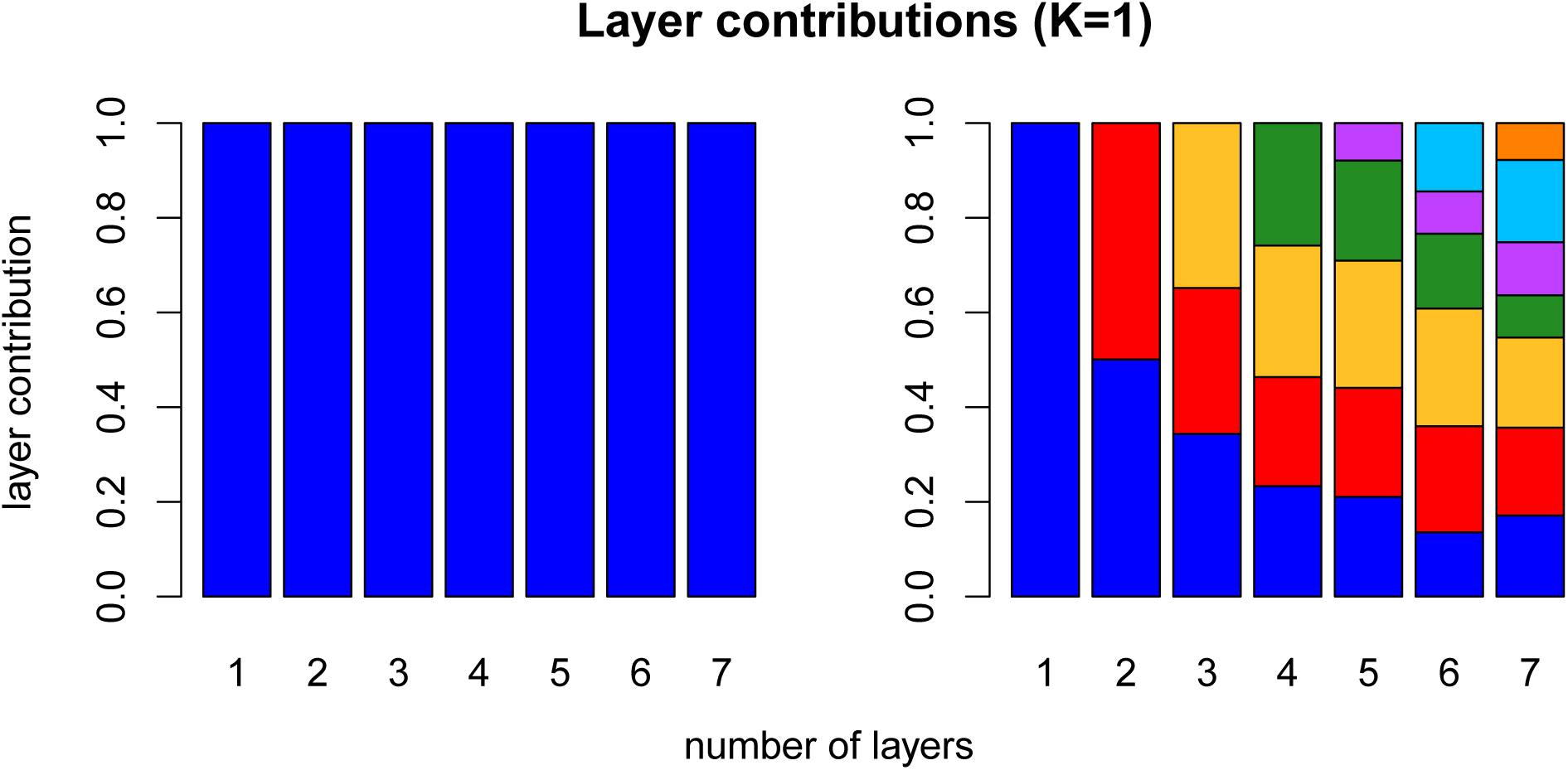
Results for data simulated using *K* = 1, showing layer/cluster contributions (i.e., how much each layer/cluster contributes to total covariance), from `conStruct` runs using *K* = 1 through 7 for the spatial model (left), and the nonspatial model (right). In each run of the nonspatial model, a single layer explained nearly all the covariance (additional bars are present but not visible).

## Empirical Applications

To further demonstrate the utility of this method, we also applied `conStruct` to empirical population genomic data from two systems: a contact zone between two poplar species in northwestern North America, and a large North American sample of black bears.

### Poplars

Trees in the genus *Populus* (poplars, aspens, and cottonwoods) are distributed throughout the Northern Hemisphere; species in the genus regularly co-occur and, where they do, they frequently hybridize [53,54].

*Populus trichocarpa*, the black cottonwood, and *Populus balsamifera*, the balsam poplar, have a broad zone of overlap in the Pacific Northwest, where they are hypothesized to hybridize [55,56]. Both species are sampled over a large geographic region, and show spatial patterns of genetic and phenotypic variation [57,58], making the system well-suited for application of our method. We organize the results of our analyses around the following questions:

1. To what degree has hybridization blurred the boundaries between *trichocarpa* and *balsamifera*? (As an extreme case, does genetic differentiation support these as separate species, as opposed to a single cline of ancestry?)
2. Does the only significant boundary of population structure fall along the species boundary (if any), or is there sub-structuring within species?
3. Does the strength of isolation by distance differ between inferred layers? This may indicate, e.g., different speeds of postglacial expansion or primary modes of dispersal.

We use data from [55], consisting of 434 individuals sampled from 35 drainages genotyped at just over 33,000 loci (map of the sampling shown in Fig S17). The number of individuals per drainage ranged between 1 and 50, with most sampling concentrated on *trichocarpa* drainages. The data were generated using RAD-seq, and showed a strong pattern of bias in allelic dropout (the majority of missing data were from drainages with only *Populus balsamifera* individuals). To ameliorate some of the problems that arise when there is a strong bias in which data are missing, we dropped loci for which any data were missing, resulting in just over 20,200 loci retained for analysis. We then analyzed these data, grouped by drainage, using both the spatial and nonspatial `conStruct` models, using *K* = 1 through 7, and compared these models using cross-validation. For comparison, we also ran fastSTRUCTURE [16] using *K* = 1 through 7. The results of all these analyses are shown in Figs 5 and 6, as well as in Figs S18 - S23 in the Supplementary Materials.

**Fig 5.**
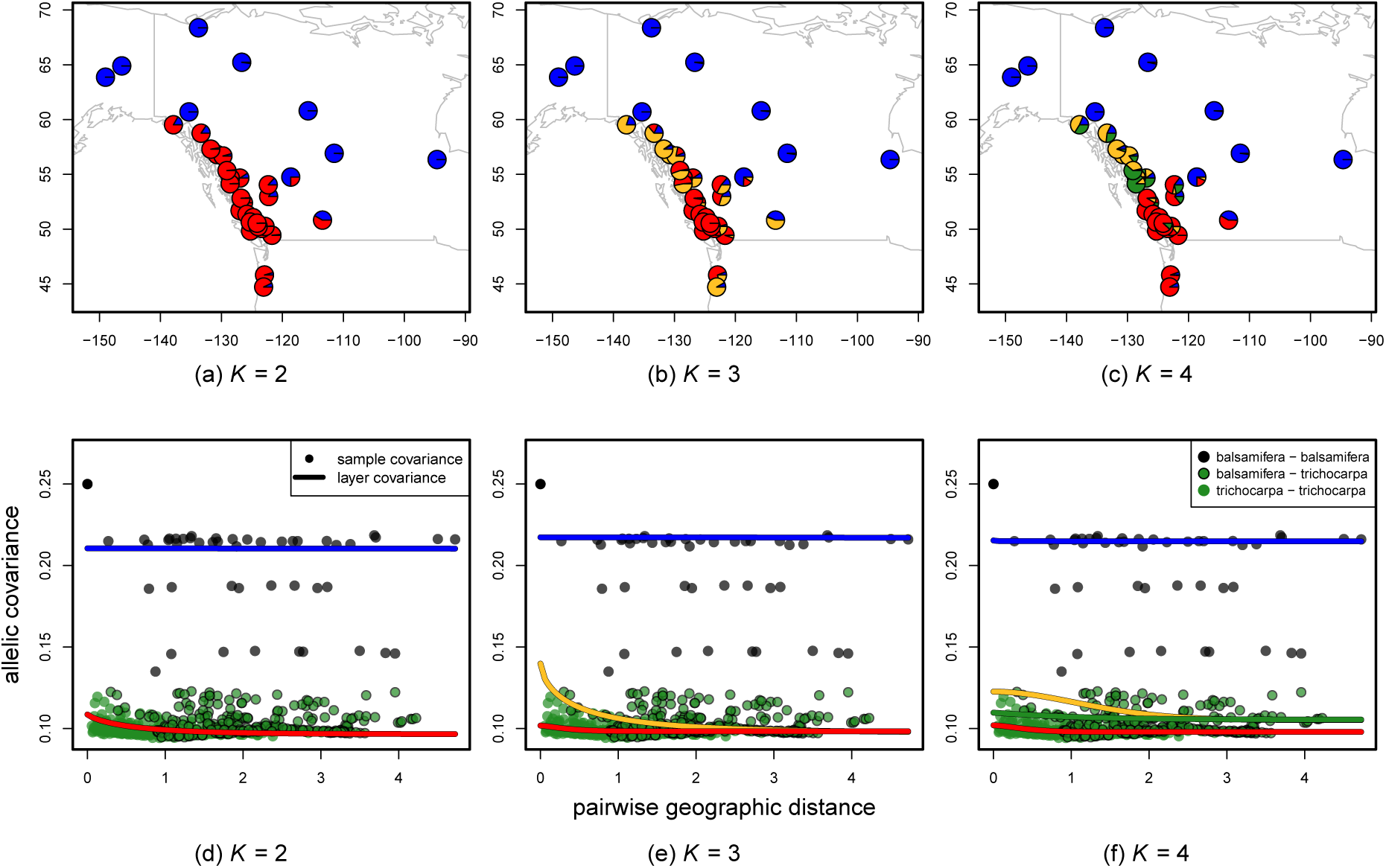
Maps of admixture proportions estimated for the *Populus* dataset using the spatial model for *K* = 2 through 4 (a-c), as well as the corresponding layer-specific covariance curves estimated under each model (d-f).

**Fig 6.**
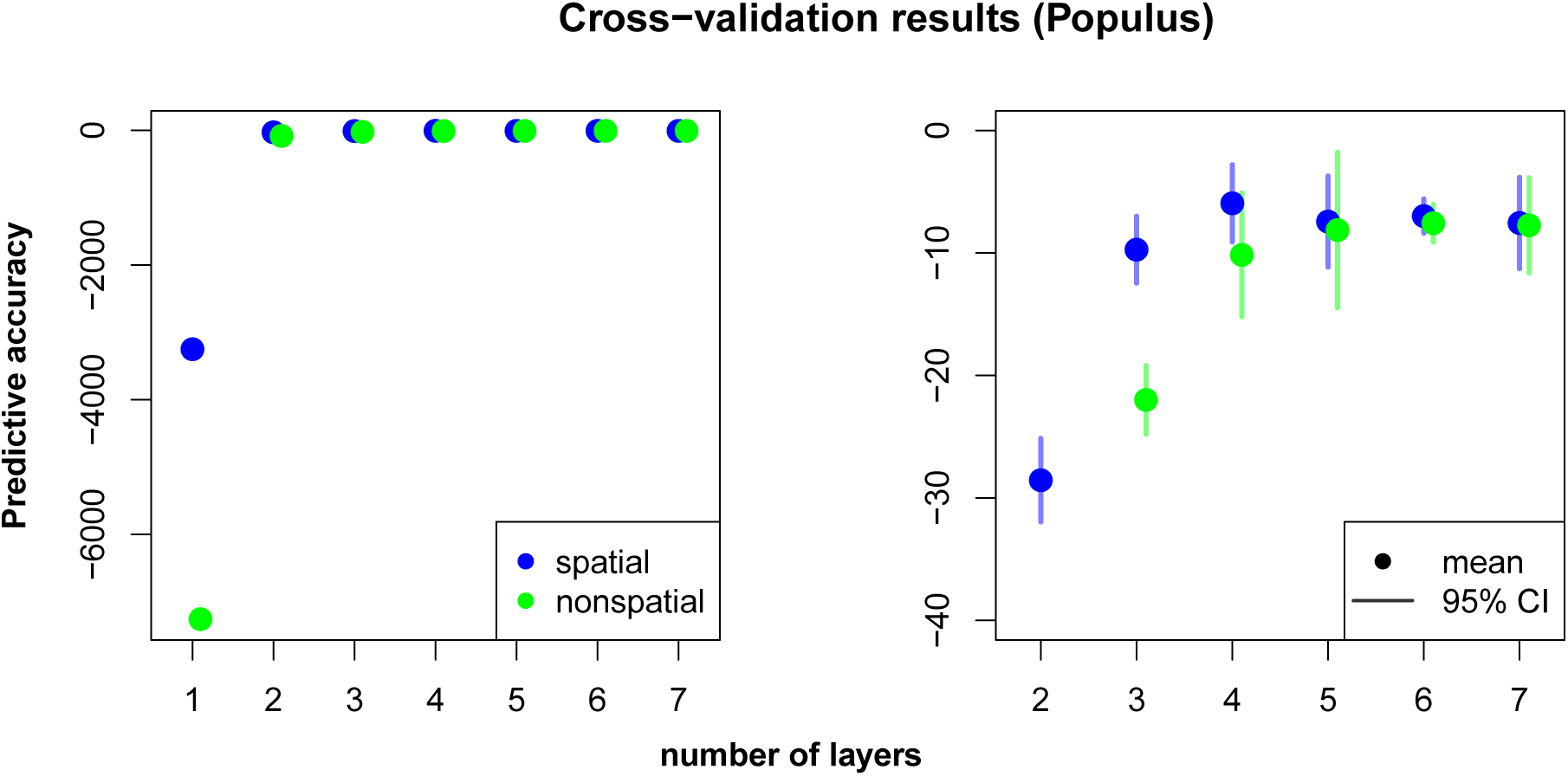
Cross-validation results for *Populus* dataset comparing the spatial and nonspatial `conStruct` models run with *K* = 1 through 7. The first panel in each row shows all results; the second panel zooms in on the results from analyses run with *K* = 2 through 7.

All models with *K* > 1 assigned the majority of each of the two species to distinct layers, with some populations drawing ancestry from multiple layers. Based on cross-validation results, we view the *K* = 3 spatial model as a sufficient description of the data, with additional structure of uncertain significance. This provides strong support for discrete population structure between the two species, with some admixture, rather than a single, continuous cline of ancestry. At all values of *K* > 1, discrete population structure was mostly partitioned along species lines; at values of *K* above 2, further discrete substructure was inferred within the *P. trichocarpa* samples, with no substructure within *balsamifera*. There was also strong support for isolation by distance in the dataset, but most of this signal seems to derive from the *P. trichocarpa* samples: as seen in Figs 5d-f and S20, there is almost no isolation by distance within the *balsamifera* layer (*α*_*D*_ ≈ 0). Both points are in agreement with [59], who found low diversity within the region’s *balsamifera*, probably as the result of a recent postglacial expansion.

A consistent split between layers within *trichocarpa* fell along the “no-cottonwood belt,” a region along the central coast of British Columbia in which black cottonwood is absent (the break between yellow and red, for *K* ≥ 3). The no-cottonwood belt is hypothesized to divide the species’ distribution into northern and southern groups, which, in a provenance test, were experimentally shown to display differences in ecologically relevant phenotypes (e.g., pathogen resistance, [60,61]). At higher values of *K*, drainages at the southern tip of *trichocarpa* sampling begin to split out into their own layers, perhaps due to introgression from the southern neighbors *P. angustifolia* or *fremontii* [55, 62]

Both nonspatial `conStruct` and fastSTRUCTURE displayed the successive partitioning of space and the clines of admixture seen in the simulation results. The details of each were somewhat different (Fig S19 vs. Fig S23), and also differed across the replicate analyses.

### Black bears

The American black bear, *Ursus americanus*, is endemic to North America and has a broad distribution across the continent. During the last glacial maximum, black bears were confined to isolated glacial refugia, from which they subsequently expanded to occupy their current range [63–66], likely leading to both continuous and discrete patterns of genetic structure. We organize our results around the following questions:

1. How many distinct populations are reflected in modern patterns of genetic variation?
2. How strong is isolation by distance within each inferred group?

Distinct popualations likely represent different glacial refugia, and differing strengths of isolation by distance might indicate different levels of habitat connectivity, dispersal behavior, or different postglacial histories.

We use data from [66], consisting of 95 individuals sampled across the United States and on the West coast of Canada, genotyped at just under 22,000 bi-allelic loci. The distribution of missing data across these individuals was uneven, with a few individuals representing most of the missing data, so we removed individuals with greater than 4% missing data, resulting in a final dataset of 78 individuals. We then analyzed these data, treating individuals as the unit of analysis, using both the spatial and nonspatial `conStruct` models with a *K* of between 1 and 7, and compared these models using cross-validation. We also ran fastSTRUCTURE [16] on the same dataset, using *K* = 2 through 4. The results of these analyses are shown in Figs 7 - 9, as well as in Figs S24 - S28 in the Supplementary Materials.

**Fig 7.**
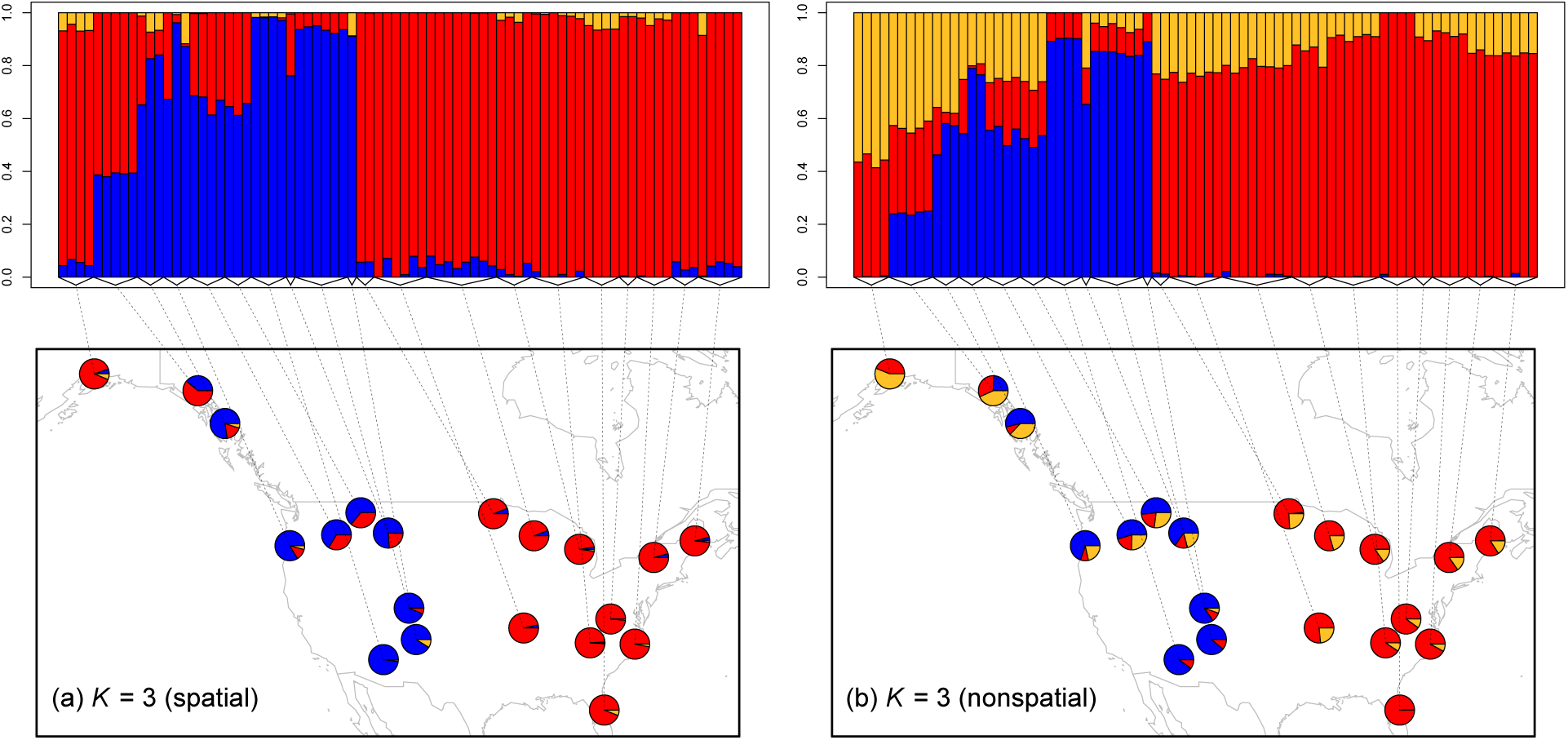
Maps of admixture proportions estimated for the black bear dataset using the spatial model (left) and the nonspatial model (right) for *K* = 3. Pies show mean admixture results across individuals within their diameter, and the admixture results for all individuals included within each group are shown in the plot above.

The results partition the sampled bears into two main groups: one (red) to the east of the Rocky Mountains, which also occurs in Alaska, the other primarily west of the Rockies (blue) (Fig 7a). The disjointed range of the blue layer likely reflects the fact that Canada was not sampled, and so the red layer may extend through the intervening (unsampled) northern Great Plains and Canadian Shield, with the blue layer presumably then stretching up into British Columbia.

Additional layers in the spatial model have strong statistical support up until around *K* = 5 or 6 (Fig 8), but additional spatial layers beyond *K* = 2 contribute little to total covariance (Fig 9). The locations of admixed individuals are consistent with a scenario of postglacial expansion from two refugia, one in the American Southwest and one in the American Southeast, meeting near the Northwest coast of North America and the Cascade Range. However, lack of any samples from Canada and Mexico, and lack of denser sampling across northern North America, make more detailed interpretations untrustworthy.

**Fig 8.**
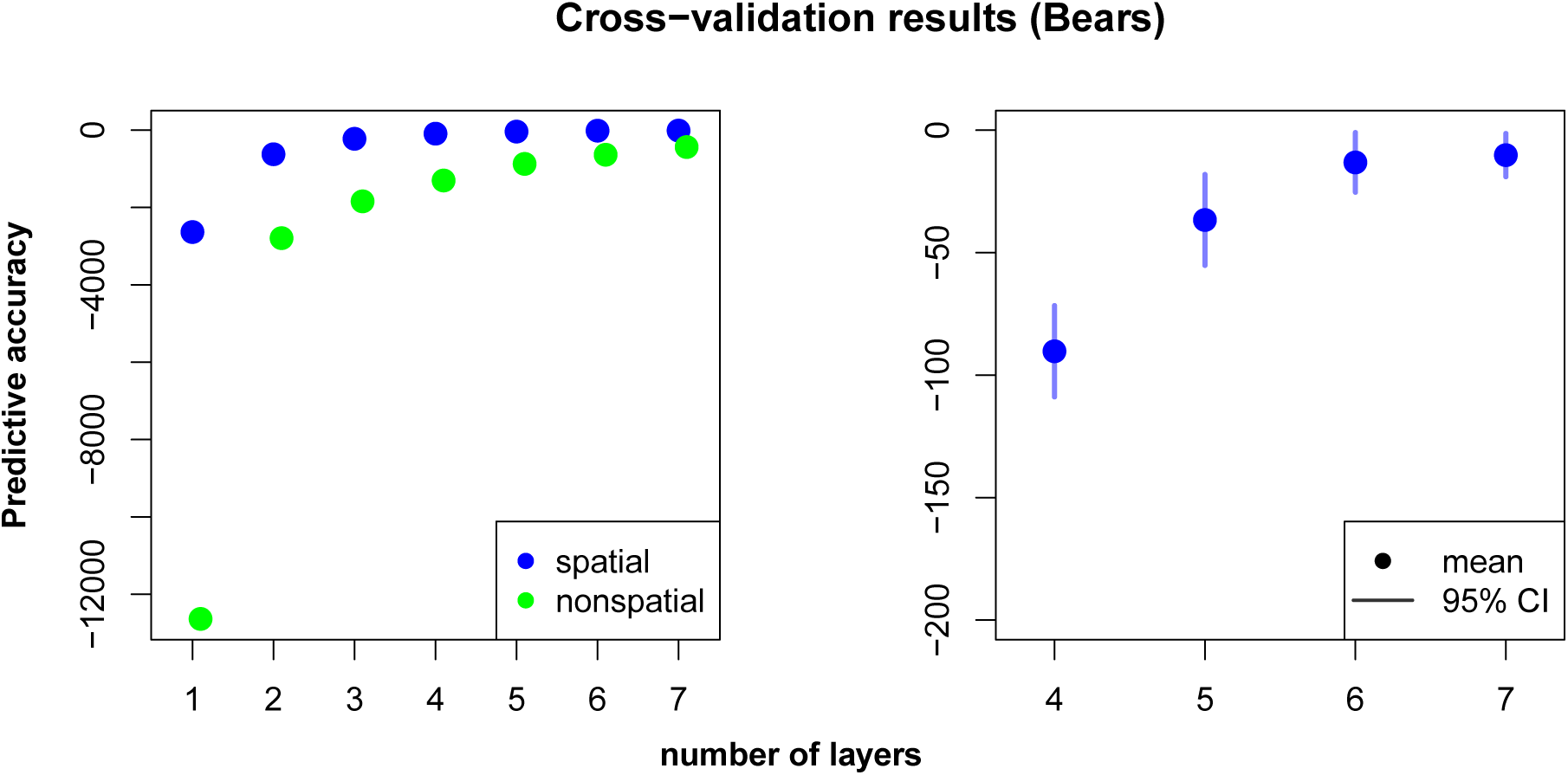
Cross-validation results for the black bear dataset, comparing spatial and nonspatial `conStruct` models run with *K* = 1 through 7. The first panel in each row shows all results; the second panel zooms in on the results from the spatial analyses run with *K* = 4 through 7.

**Fig 9.**
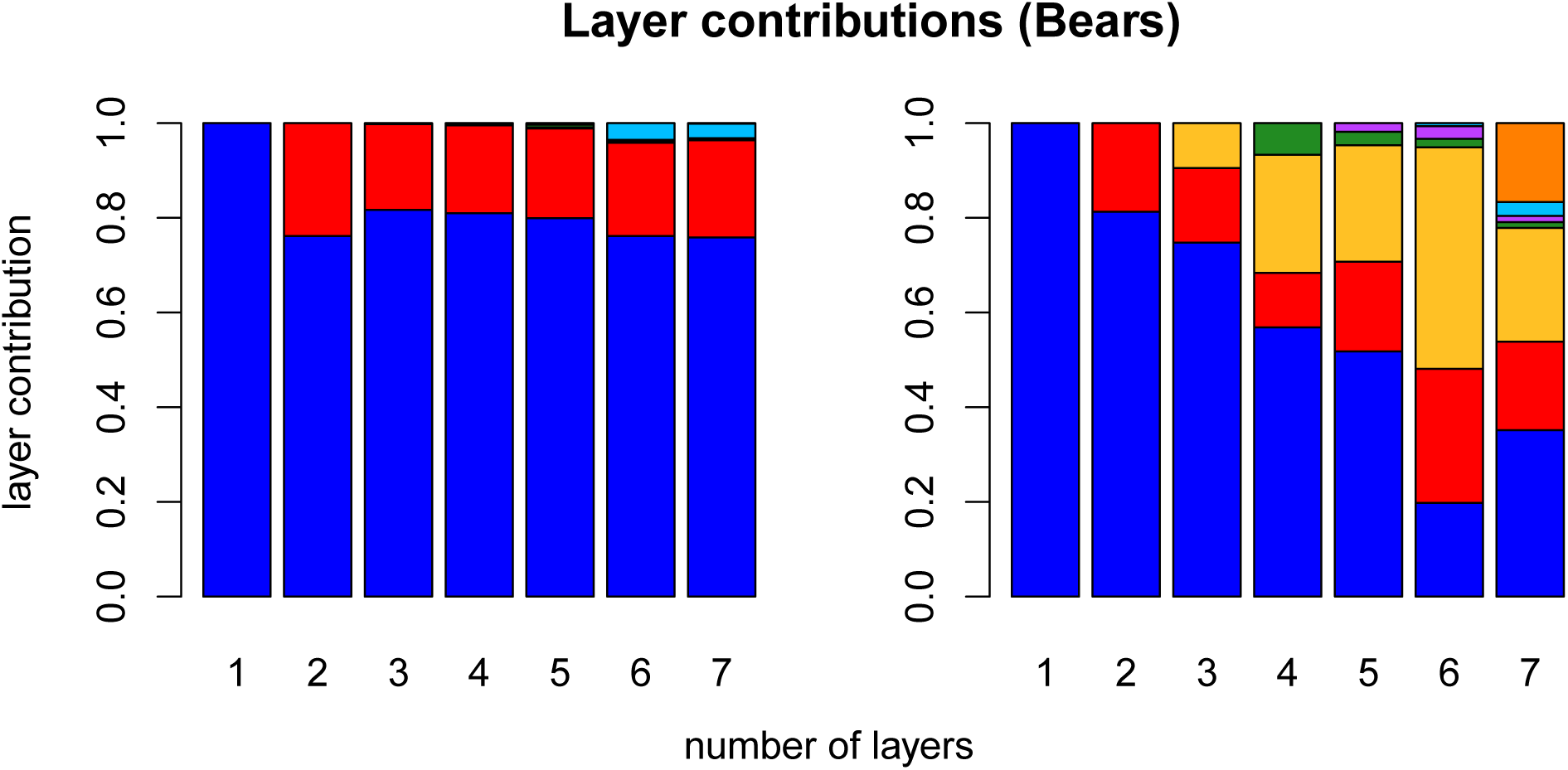
Layer/cluster contributions (i.e., how much total covariance is contributed by each layer/cluster), for all layers estimated in runs using *K* = 1 through 7 for the spatial model (left), and for all clusters using the nonspatial model (right). For each value of *K* along the x-axis, there are an equal number of contributions plotted. Colors are consistent with Fig 7.

Results from the nonspatial model clearly exhibit the tendency of nonspatial clustering algorithms to describe continuous spatial patterns of divergence using gradients of admixture between clusters. For example, in Fig 7b, the third cluster (in gold) exhibits a clear East-West gradient that overlays the discrete structure between the Southwest cluster and the Southeast. The results from fastSTRUCTURE are are not identical to those obtained using the nonspatial model, but they do show the same tendency: e.g., at *K* = 3, fastSTRUCTURE splits the westernmost Alaskan samples out of the cluster with the eastern samples, and at *K* = 4, it splits them into their own cluster entirely (Fig S28).

Across all values of *K* for which we ran `conStruct`, we see strong support for the spatial model over the nonspatial model (Fig 8). This pattern may resolve a discrepancy between our results and previous analyses that split Alaskan and British Columbian bears out into their own cluster with an inferred Beringian glacial refugium [64–66]. Our model, which explicitly incorporates a spatial decay of relatedness, allows somewhat genetically differentiated individuals that are sampled far from one another to belong to the same layer, instead of splitting these individuals out into successive clusters (e.g., Fig S24d vs S25d).

## Discussion

In this paper, we have presented a statistical framework, `conStruct`, for simultaneously modeling continuous and discrete patterns of population structure. By employing the sensible default assumption that relatedness ought to decay with geographic distance, even within a population, we avoid erroneously ascribing population differentiation to discrete population clusters. To aid comparison between models, we present a cross-validation approach as well as a way to describe the contribution of each spatial layer to the model (but caution against overly strict interpretation of either).

The method performs well on simulated data: we accurately infer the admixture proportions used to simulate the data and accurately pick the simulating model as the best model using our cross-validation procedure. Two empirical applications of `conStruct` to samples of North American poplars and black bears yield reasonable results, and demonstrate that, by acknowledging isolation by distance, real datasets can be better described using fewer layers.

The proposed method combines the utility of model-based clustering algorithms with a biologically realistic model of isolation by distance. We anticipate that `conStruct` will be useful for identifying populations and determining samples’ ancestry in and across them, especially when the populations exhibit spatial patterns of relatedness.

### Comparison to nonspatial model-based clustering

Above, we showed that (a) the nonspatial `conStruct` model recapitulates results of other, commonly-used nonspatial clustering methods, and (b) `conStruct` can concisely capture spatial structure, which will be common within populations. Given this, when should methods without spatial capability be used? One advantage these have over `conStruct` is speed when the number of samples is large. Although `conStruct`’s computation time is independent of the number of loci included in the dataset (after the initial calculation of the allelic covariance), it currently scales poorly with number of samples. The computationally limiting step is the inversion of the parametric covariance, which scales more than quadratically with the number of samples, whereas computation time for, e.g., STRUCTURE, scales linearly with number of samples. Our speed, on datasets with large sample sizes, could be improved by adopting rank-one updates to the inverse of the covariance matrix (e.g., [67, 68]) when updating a sample’s admixture proportions, which alters only a single row/column of the covariance matrix. We have not implemented this yet, as it would likely mean losing the ability to do efficient, Hamiltonian Monte Carlo sampling of our parameters.

For a relatively small number of samples, `conStruct` can be much faster than other nonspatial clustering methods. On a desktop machine, using a single 4.2 GHz Intel Core i7 processor, an analysis of the black bear dataset (78 samples, 21,000 loci) running `conStruct`’s spatial model with 4 layers for 5,000 MCMC iterations (which was more than sufficient for convergence) took 2.8 hours. On the same machine and dataset, a fastSTRUCTURE run with *K* = 4 took 90.2 hours. However, for a large number of samples, fastSTRUCTURE (or ADMIXTURE) would be much faster.

### Choosing the “best” number of layers

Although we recognize the utility of choosing a single, “best” value of *K*, and using only that analysis to communicate results, we emphasize that the choice of best *K* is always relative to the data in hand and the questions to be answered. From a statistical perspective, unless the data were generated under the model itself, the support for larger values of *K* is likely to increase with increasing amounts of data. In the limit of infinite data, the best value of *K* may be the number of samples included in the dataset [69].

From a biological perspective, it is important to stress that patterns of relatedness between individuals and populations are shaped by complex spatial and hierarchical processes. All individuals within a species (and indeed, all individuals across all species), are related to one another in some way, and summarizing that relatedness with a single value of *K* may be reductive or misleading. We therefore encourage users to perform analyses across different values of *K* and observe which layers split out at what levels (this is conceptually similar to taking successively shallower cross-sections of the population phylogeny), and also to take the results of the proposed cross-validation procedure with a large grain of salt. Calculating layer contributions may also be a useful heuristic, as it can reveal layers with statistical support but small biological import.

Although we believe our model adds spatial realism to the groups used by clustering methods, it is important to note that the layers detected by our method do not necessarily correspond to distinct, ancestral populations; nor does a non-zero admixture proportion indicate that admixture (i.e. gene flow) must have occurred. Both groupings and admixture proportions should be viewed as hypotheses that should be subject to further testing (see [70] for an in-depth discussion of these points).

### Implications for management and conservation

Because isolation by distance is common, a likely result of applying `conStruct` to existing data is that some populations previously identified using nonspatial clustering methods may be collapsed into each other. This “lumping” might better reflect biological reality, but may also have implications for management decisions and conservation policy, both of which are often predicated on the identification of discrete “management units” (MUs) identified using genetic data [2–4].

It is therefore important to stress that individuals sampled from the same `conStruct` layer may be quite genetically diverged from one another, perhaps especially at loci underlying adaptive traits, and that a `conStruct` layer may still contain multiple distinct MUs worthy of independent protections. Alternatively, the inclusion of multiple MUs into a single `conStruct` layer may occur if these populations are currently (or were recently) exchanging migrants, and thus might emphasize the importance of maintaining habitat corridors between demes, or of implementing an integrated conservation plan across multiple demes within a layer.

### Caveats and considerations

There are a few important caveats to consider in the interpretation of `conStruct` results. First, we have modeled allelic covariance within a layer as a spatial process. Although there is flexibility built into the model about the shape of that covariance, inference may be misleading if the sampling geography departs radically from the way the sampled organisms disperse (or have dispersed) on their landscape. For example, if we were to run a `conStruct` analysis using geographic distances between sampled individuals of greenish warblers [71] or *Ensatina* salamanders [72] — two canonical examples of rings species — we might get misleading results. This is because distance between locations on either side of the species’ distributions (across the Tibetan plateau and the Central Valley, respectively) is not representative of the path traversed in the coalescent of a pair of alleles sampled at those locations.

A second caveat is that, in some instances, membership in the same layer may not mean that samples are particularly related. If covariance within a layer decays sharply with distance, and the layer-specific relatedness parameter *ϕ*^(*k*)^ is low, individuals separated by a large spatial distance may be in the same layer but have very low pairwise relatedness. It is possible that this is happening in Fig S18. At *K* = 3, the southernmost populations of *P. trichocarpa* are in the gold layer, whose other neighbors are to the north, with an intervening group of populations in the red layer, and at *K* = 5, those southernmost samples split out and become their own layer. Again, we encourage users to run analyses across multiple values of *K*, and to examine the spatial covariance functions within layers when interpreting results.

### Extensions and future directions

There are several ways in which the model described in this paper might be extended or improved. For example, we currently assume that all layers within a model are equally unrelated (a star population phylogeny, although the branches can have different lengths thanks to the *ϕ*^(*k*)^ parameter), similar to the F-model of [12]. However, we could extend the existing model by implementing a relatedness structure between the layers by, for example, estimating a population phylogeny between them (e.g, [6]).

In addition, here we have assumed that samples have known geographic coordinates, and that they draw ancestry from layers only at those sampled locations. A natural extension would be to attempt to “geo-locate” the ancestry of samples without geographic coordinates [35]. We could also imagine letting samples draw ancestry from other geographic coordinates, as we have done in a previous approach [37] to model long distance dispersal. We could even allow entire layers to bud off of a particular location on another layer. This would enable more explicit modeling of range expansion or domestication, in which a set of individuals are thought to have ancestry that originated from a particular geographic location embedded in a larger pattern of isolation by distance.

A final direction would be to model relatedness within a layer as a spatiotemporal process, in which covariance decays both with distance in space and in time. As the number of genotyped historical or ancient samples increases, it is becoming possible to ask whether there is genetic continuity at a point in space across time, or whether populations are being replaced [73–77]. However, we expect allele frequencies to change through time in a population, even without replacement, simply due to drift. Therefore, a natural way to test for population replacement is to estimate the rates at which relatedness within a layer decays with time in the same way we do in the current model with space, in which case a change in discrete population structure across space is comparable to population replacement across time.

## Materials and Methods

### Model rationale

#### Drift, admixture, and space

Here we sketch a simple model of allele frequencies and their covariances, to justify the form given in the main text.

#### Drift

We first provide a simple model of allele frequencies within a layer. Imagine a sample *i* that draws all of its ancestry from layer *k*. The allele frequency in sample *i* at locus *l*, denoted *F*_*i,l*_, can be written as the sum

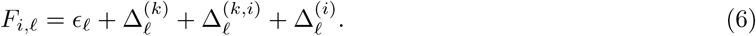

The first term is the ancestral allele frequency *ϵ*_*l*_ shared by all samples; the second is the deviation from that ancestral frequency due to drift in the ancestral population of the *k*th layer, which is shared by all samples within the layer. The third term is the deviation of the *i*th sample away from the *k*th layer mean due to the spatial process of drift and migration within the layer. The final term is the deviation specific to the *i*th sample, which captures drift not shared by all samples at the population level (i.e., subpopulation-specific drift due to, e.g., inbreeding). We will assume that these four deviations are all uncorrelated with each other.

If we have two samples *i* and *j* drawn from layer *k*, their covariance across loci will be

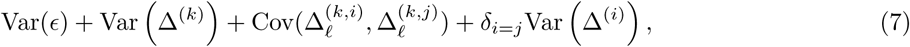

where the quantity *δ*_*i*=*j*_ is an indicator variable that equals 1 when *i* is equal to *j* and 0 otherwise, as in Eq. (3).

#### Admixture

The model above describes the simple case in which samples draw 100% of their ancestry from only a single layer each. To accommodate admixture between layers, we model sampled genomes as drawn from allele frequencies that are weighted averages of the local frequencies in each layer from which they draw ancestry. The weights, 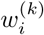, describe the “admixture proportion” of sample *i* in layer *k*. (Note that 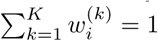 for each *i*) These can be interpreted as the proportion of the genome in the *i*th sample that came from the *k*th layer (or the probability that an allele at a locus is drawn from layer *k*). The allele frequency in the *i*th sample at the *l*th locus can therefore be written as:

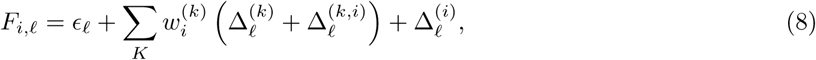

and so the covariance between *i* and *j* across loci is

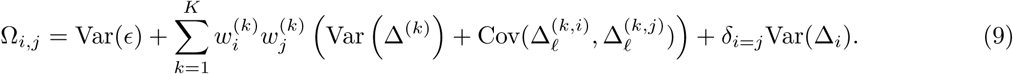

#### Space

Under our nonspatial model, we assume that 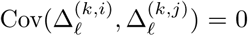, so that the only additional covariance between *i* and *j* (above that induced by a shared ancestral frequency at each locus) is due to the drift in the ancestral population of their layer (the variance of which is *ϕ*^(*k*)^). Under our spatial model we assume that some of the covariance in allele frequencies between *i* and *j* decays as a function of the geographic distance between the pair, *D*_*i,j*_, so that

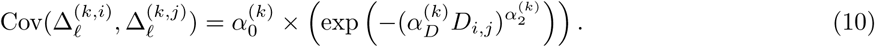

We note that this form is chosen for its flexibility, and not because it matches any explicit population genetic model of isolation by distance.

### Allelic covariance

To see how to arrive at our empirical variance-covariance expression (Eq. (1)) pick a random locus and let *A* and *B* be randomly drawn alleles at that locus from populations *i* and *j* respectively. If *i* = *j*, then these are chosen *without* replacement. Suppose these are each coded as ‘0’ or ‘1’ (where ‘0’ denotes a reference allele), but we randomly “flip” this coding, so that we let *X* = *A* and *Y* = *B* with probability 1/2, but otherwise we let *X* = 1 – *A* and *Y* = 1 – *B*. We then let 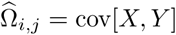. Sampling without replacement makes the statistic insensitive to sample size, and random allele flipping makes it independent of the choice of reference allele. By conditioning on the flip, and using the fact that 𝔼[*X*] = 𝔼[*Y*] = 1/2, Eq. (1) comes from the observation that

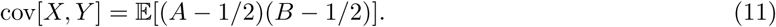

The factor of *δ*_*i,j*_ /(*N*_*i*_ – 1) comes from the fact that if *i* = *j*, then 𝔼[*AB*] is the probability that both alleles drawn without replacement are reference, which is 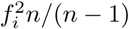.

#### Likelihood

If allele frequency deviations are well approximated by a Gaussian, their sample allelic covariance is a sufficient statistic, so that calculating the likelihood of their sample allelic covariance is the same as calculating the probability of the frequency data up to a constant. We can therefore model the covariance of the sample allele frequencies, 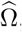, as a draw from a Wishart distribution with degrees of freedom equal to the number of loci *L* across which the sample allelic covariance is calculated:

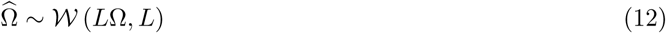

where 𝒲 is the Wishart likelihood function.

A benefit of directly modeling the sample allelic covariance is that, after the initial calculation of the sample covariance matrix, the computation time of the likelihood is not a function of the number of loci, so inference can theoretically be done using whole genome data.

### Models, parameters, and priors

#### Spatial versus nonspatial

In this paper, we discuss two types of models, spatial and nonspatial, each of which can be implemented with different numbers of layers/clusters. The spatial model is parameterized as in Eq. (9), and the nonspatial model is a special case of the spatial model with all *α* parameters is set to 0. The nonspatial model therefore has 3*K* fewer parameters than the spatial model, because there are three *α* parameters that describe the continuous differentiation effect of distance in each layer.

#### Single layer

Each of these models can be run with a single layer (*K* = 1), in which case the layer-specific covariance parameter *ϕ*^(*k*)^ and the global covariance parameter γ become redundant. The single-layer model is therefore a special case of the multi-layer model, in which we set *ϕ* to zero. For the spatial model, the single-layer parametric covariance is:

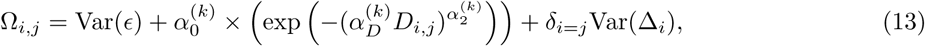

and for the nonspatial model, it is:

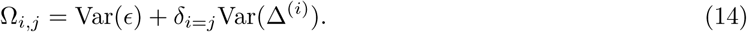

#### Priors

We use a Bayesian approach to parameter inference. A table of all parameters, their descriptions, and their priors is given in Table 1.

### Cross validation

We employ Monte Carlo cross-validation approach for model comparison [78]. This procedure generates a mean predictive accuracy for each model and each value of *K*, as well as a confidence interval around that mean, which can then be used for model comparison or selection. Briefly, we follow the following procedure:

1. For each of *X* replicates:

a. partition the allele frequency data into a 90% “training” partition 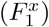 and a 10 %“testing” partition 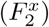
b. run our inference procedure on the training partition to estimate model parameters *θ*_*mk*_ for:

i. *m*: the spatial and the nonspatial model
ii. *k*: the number of layers/clusters 1 through *K*
c. calculate the mean log likelihood of the testing data partition over the posterior distribution of training-estimated parameters for each model (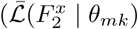, henceforth 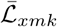)
d. generate standardized mean log likelihoods, *Ƶ*_*xmk*_, across models:

i. identify the highest mean log likelihood, 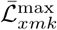
ii. subtract 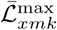 from 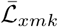 across all models, such that the standardized log likelihood, *Ƶ*_*xmk*_, of the best model is 0, and less than 0 for all inferior models.
2. For each model (i.e., each combination of *m* and *k*) calculate the mean 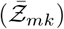 standardized log likelihood of the testing data partition across *X* replicates, as well its standard error 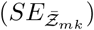 and 95% confidence interval 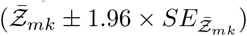.

### Calculating layer contributions

Let *A* and *B* be randomly chosen alleles from samples *i* and *j* respectively, at a randomly chosen locus (if the two populations are the same, then choose without replacement). Then, if we let *U* = 2(*A* – 1/2) and *V* = 2(*B* – 1/2), since *U* and *V* take the values ±1, so as in Eq. (11),

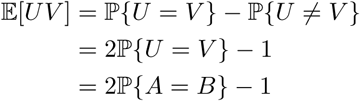

To translate, ℙ{*U* ≠ *V*} is the probability that the alleles from our two focal samples agree with each other, while ℙ{*U* ≠ *V*} is the probability that they disagree. This implies that 𝔼[*UV*] = 1 — 2π_*ij*_, where π_*ij*_ is the probability that two randomly chosen alleles differ, which is the genetic divergence.

Now, here is a generative model that gives us the form of the covariance we have postulated. To decide whether or not *A* and *B* will agree, first each sample randomly chooses a layer: call these layers *I* and *J*. The probability that *A* chooses layer *k* is 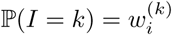, the *i*th sample’s admixture proportion in the *k*th layer. The same holds true for *B*. If they do not choose the same layer, the probability that they agree is *p*_γ_. If they do choose the same layer, then they agree with a probability 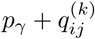 that depends on their distance apart. By the above, the probability of agreement is ℙ{*A* = *B*} = 2*cov*[*A*, *B*] *+* 1/2, and so

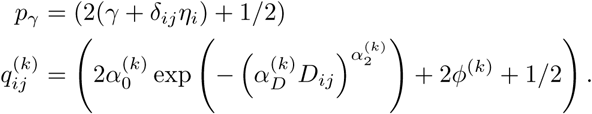

One way to summarize the contribution of each layer is to partition the probability of agreement into contributions due to agreement “in” each layer. So, the contribution from layer *k* to agreement between *i* and *j* is 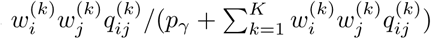 which is the probability, given that they agree, that they agree thanks to layer *k*. Similarly, the “background” contribution is 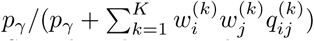. Because our signal comes from *variation* in covariance, we omit the *p*_γ_ terms. Stated in this way, this quantity is the relative contribution of the *k*th layer to the (model-based) kinship coefficient between *i* and *j*.

This suggests defining the overall contribution of layer *k* to agreement, ξ^(*k*)^ to be the average of that quantity over *i* and *j*:

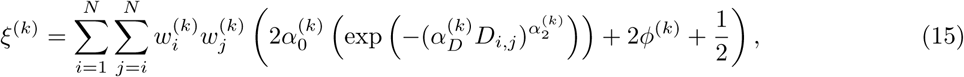

which is that layer’s contribution to agreement between samples summed over the upper triangle (excluding the diagonal) of the covariance matrix. We define the contribution of the *k*th layer, 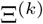, as the relative contribution of the *k*th layer to total agreement:

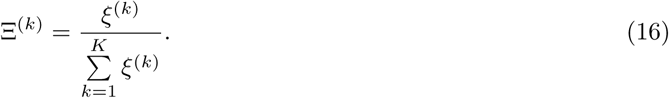

This is the quantity that is plotted in, e.g., Figures 4 and 9.

### Simulation details

We wished to simulate data under a model that had some biological realism, but at the same time had unambiguous true admixture proportions (so as to test the behavior of the method). This second requirement precluded scenarios of, e.g, recent secondary contact between populations expanding out of different refugia, which would have more biological realism, but no unambiguous ancestry proportions for admixed populations. Here, we describe in more detail the procedure we use to simulate our test dataset, using a cartoon schematic with *K* = 2 as an example (Fig 10).

Using the program ms [52], we generated discrete population structure by simulating *K* distinct populations, each of which split from a common ancestor *τ*_s_ units of coalescent time in the past, without subsequent migration between them. Then, to generate continuous differentiation within each population, at time *τ*_e_ in the past, each of these discrete populations instantaneously colonizes an independent lattice of demes, for which we use a stepping stone model with symmetric migration to nearest neighbors (eight neighbors, including diagonals).

Finally, at time *τ*_a_ in the past we generate a single dataset by collapsing those *K* discrete lattices into a single grid of demes that are admixed to various degrees from these different layers. We wish to simulate realistic patterns of admixture (and thereby set a more difficult test for the method), by generating spatially autocorrelated admixture proportions in each diverged population. To do so, we first place *K* equidistant points on the circle centered on our lattice. These points serve as “foci” of ancestry in each of the *K* layers. We then calculate the distance from each deme in the sampled lattice to each of these *K* foci, and draw admixture proportions for each deme from a Dirichlet distribution for which the concentration parameter for deme *i* in layer *k* is inversely proportional to the distance between deme *i* and focus *k*. This creates a pattern in which the admixture proportions in a given layer decreases with the distance from that layer’s focus, as might be expected if a spatial process were mediating admixture between diverged populations.

## Acknowledgements

We thank Marjorie Weber, Yaniv Brandvain, William Wetzel, Mariah Meek, Doc Edge, and Matthew Stephens for invaluable comments on the method and manuscript, as well as Quentin Cronk, who provided input on the *Populus* analyses, and Emily Puckett, who provided input on the black bear analyses. We also thank the attendees at the 2017 SSE Meeting in Portland, OR, whose votes determined the name of the method. This work was supported in part by the National Science Foundation under award number NSF #1262645 (DBI) to PR and GC, the National Institute of General Medical Sciences of the National Institutes of Health under award numbers NIH R01-GM108779 to GC, and the National Science Foundation under award numbers NSF #1148897 and #1402725 to GB.

## Supplementary Information

**Fig S1.**
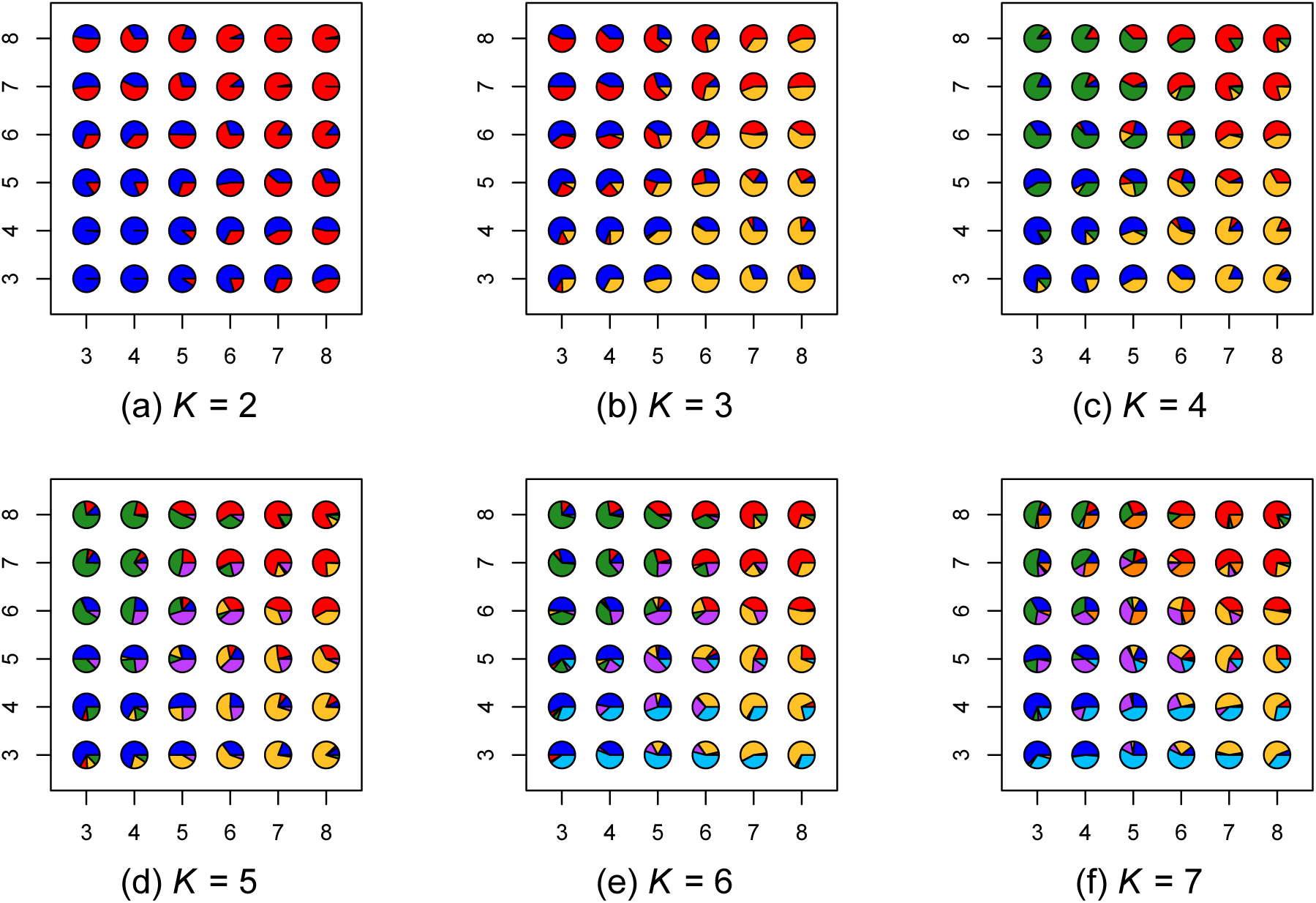
Map of admixture proportions estimated using a nonspatial model for *K* = 2 through 7. The data were simulated using one layer with nearest-neighbor symmetric migration between demes.

**Fig S2.**
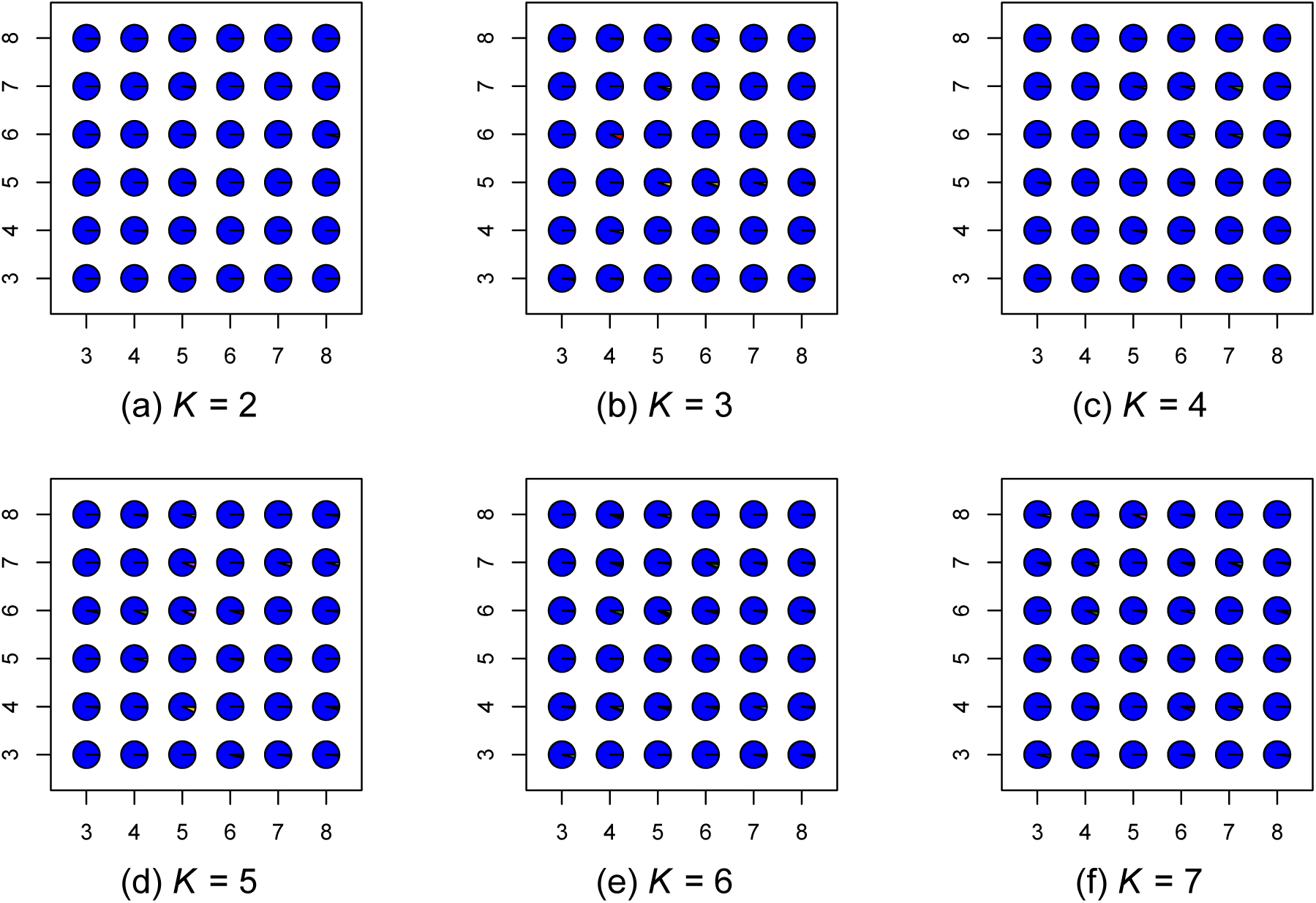
Map of admixture proportions estimated using a spatial model for *K* = 2 through 7. The data were simulated using one layer with nearest-neighbor symmetric migration between demes.

**Fig S3.**
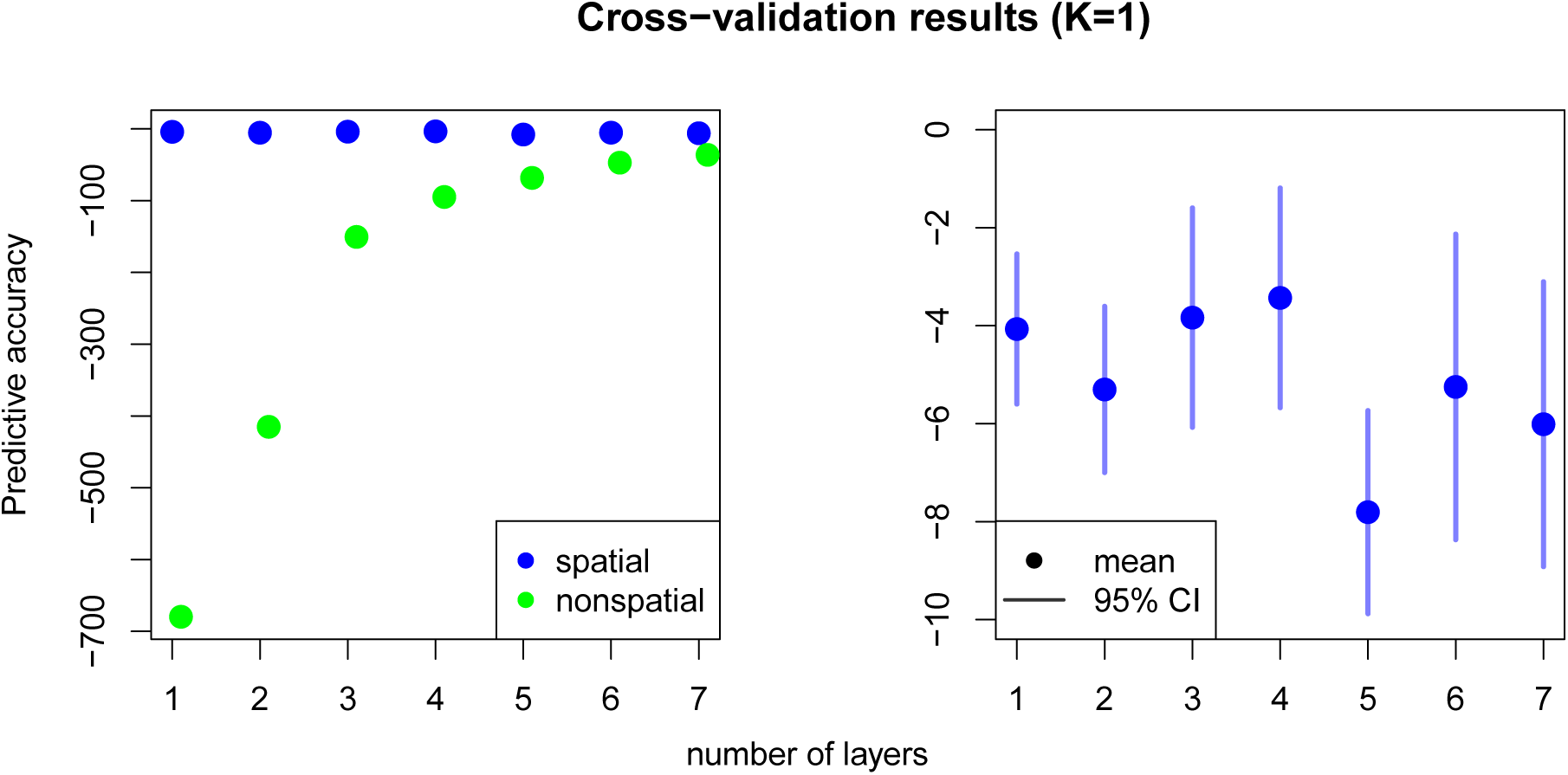
Cross-validation results for data simulated under *K* = 1, comparing the spatial and nonspatial `conStruct` models run with *K* = 1 through 7. The right panel zooms in on just the spatial cross-validation results.

**Fig S4.**
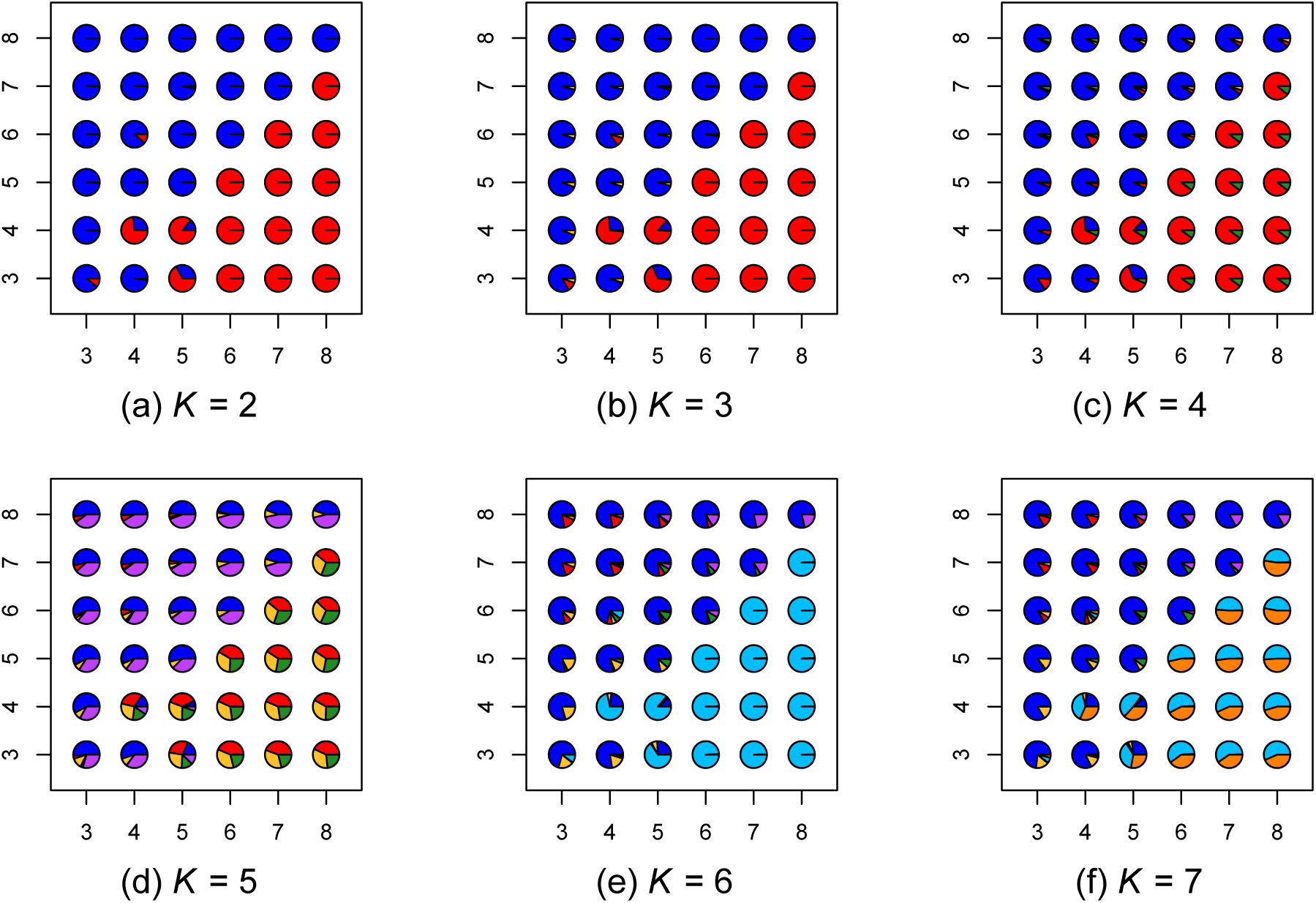
Map of admixture proportions estimated using a nonspatial model for *K* = 2 through 7. The data were simulated using two layers with nearest-neighbor symmetric migration between demes.

**Fig S5.**
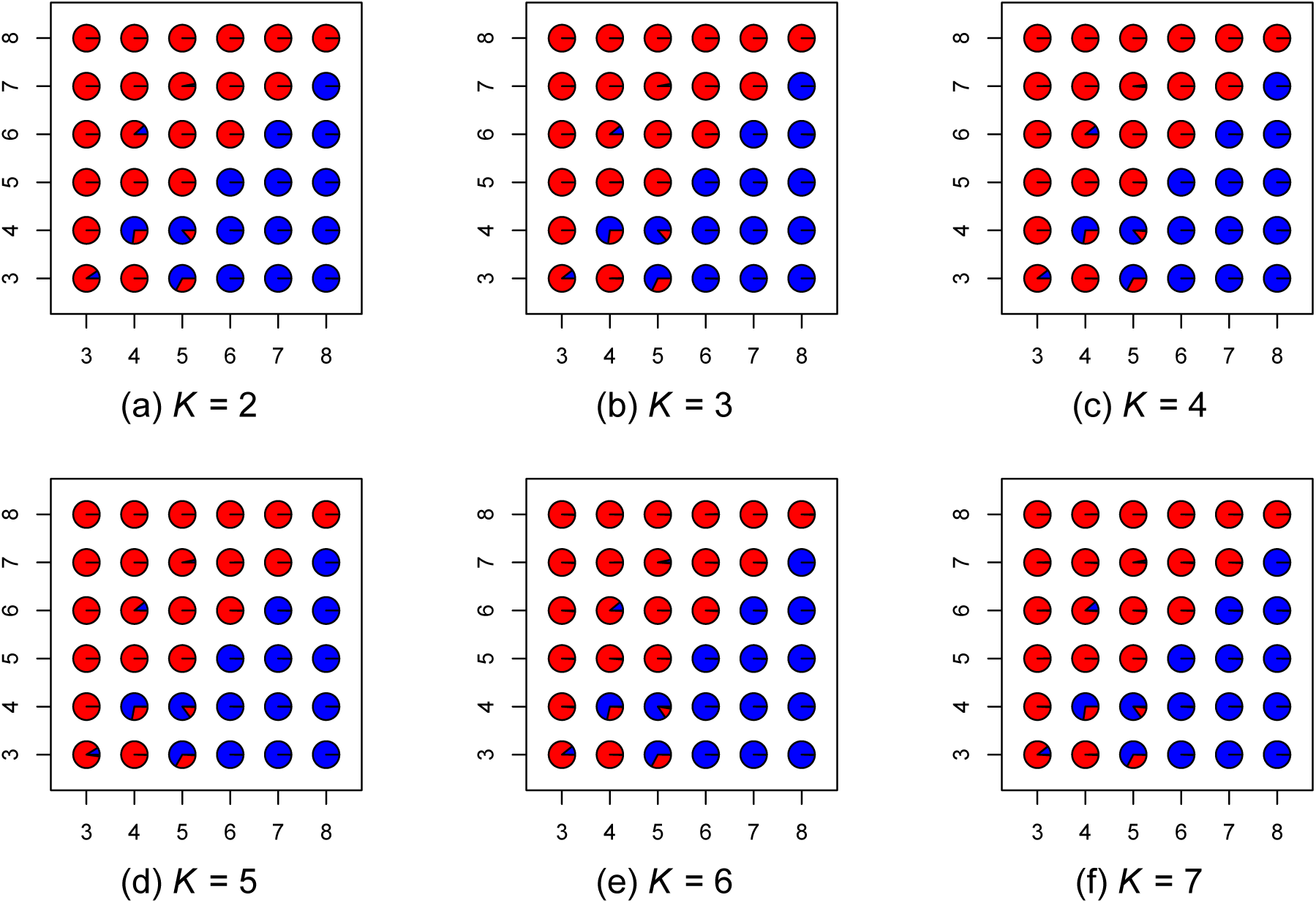
Map of admixture proportions estimated using a spatial model for *K* = 2 through 7. The data were simulated using two layers with nearest-neighbor symmetric migration between demes.

**Fig S6.**
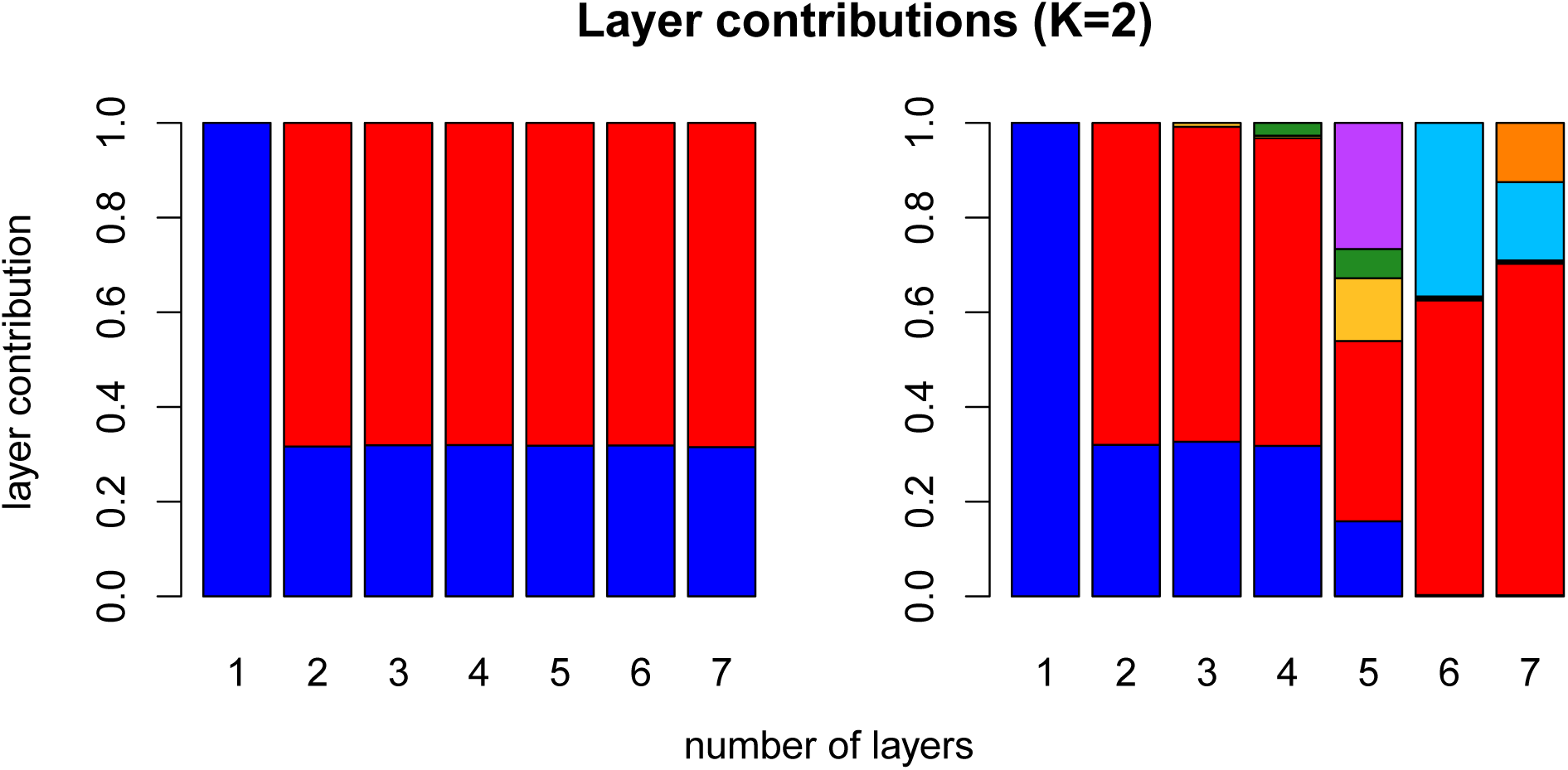
Layer/cluster contributions (i.e., how much total covariance is contributed by each layer/cluster), for all layers estimated in runs using *K* = 1 through 7 for the spatial model (left), and for all clusters using the nonspatial model (right). Data were simulated using *K* = 2. For each value of *K* along the x-axis, there are an equal number of contributions plotted.

**Fig S7.**
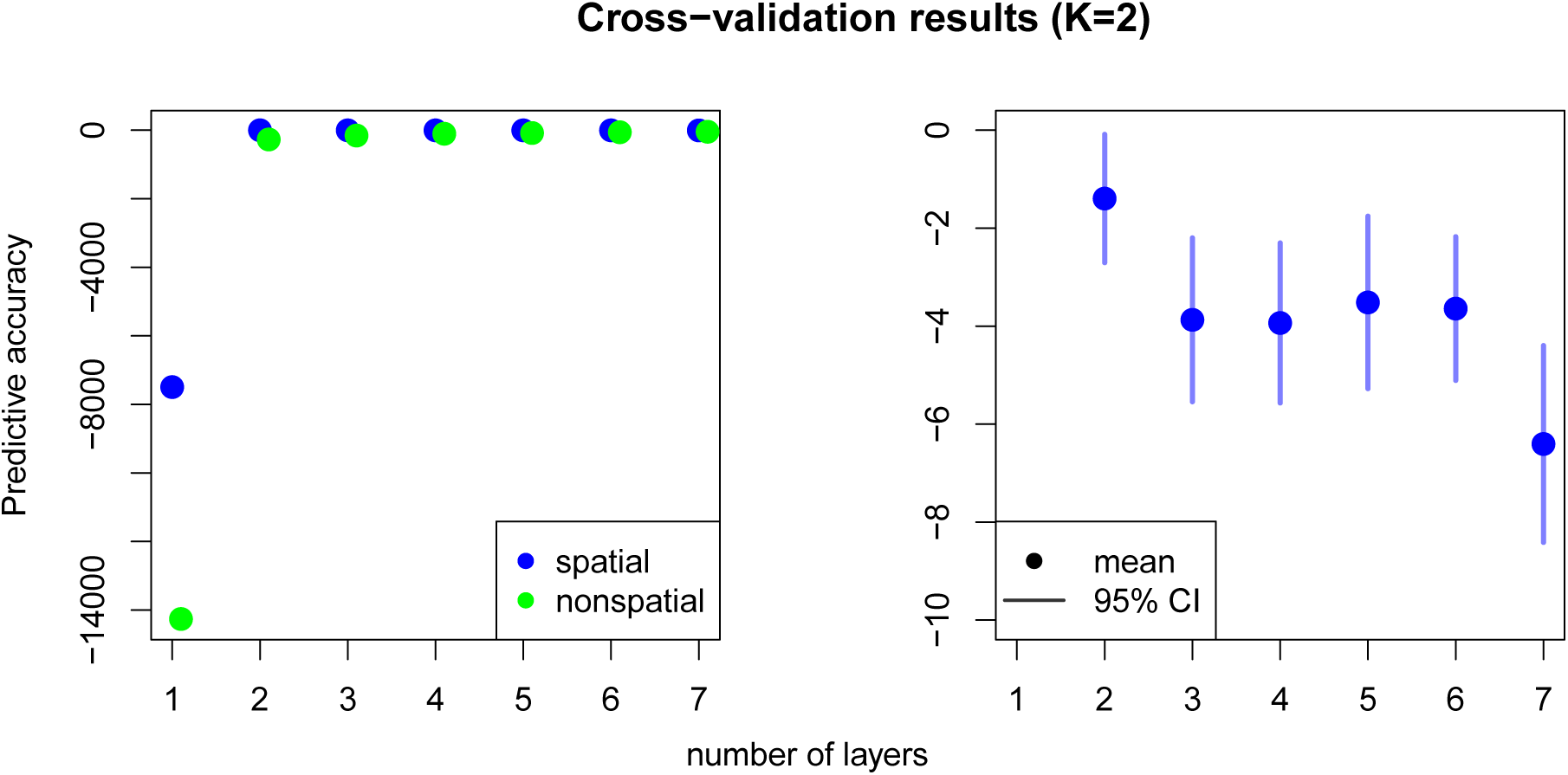
Cross-validation results for data simulated under *K* = 2, comparing the spatial and nonspatial `conStruct` models run with *K* = 1 through 7. The right panel zooms in on just the spatial cross-validation results.

**Fig S8.**
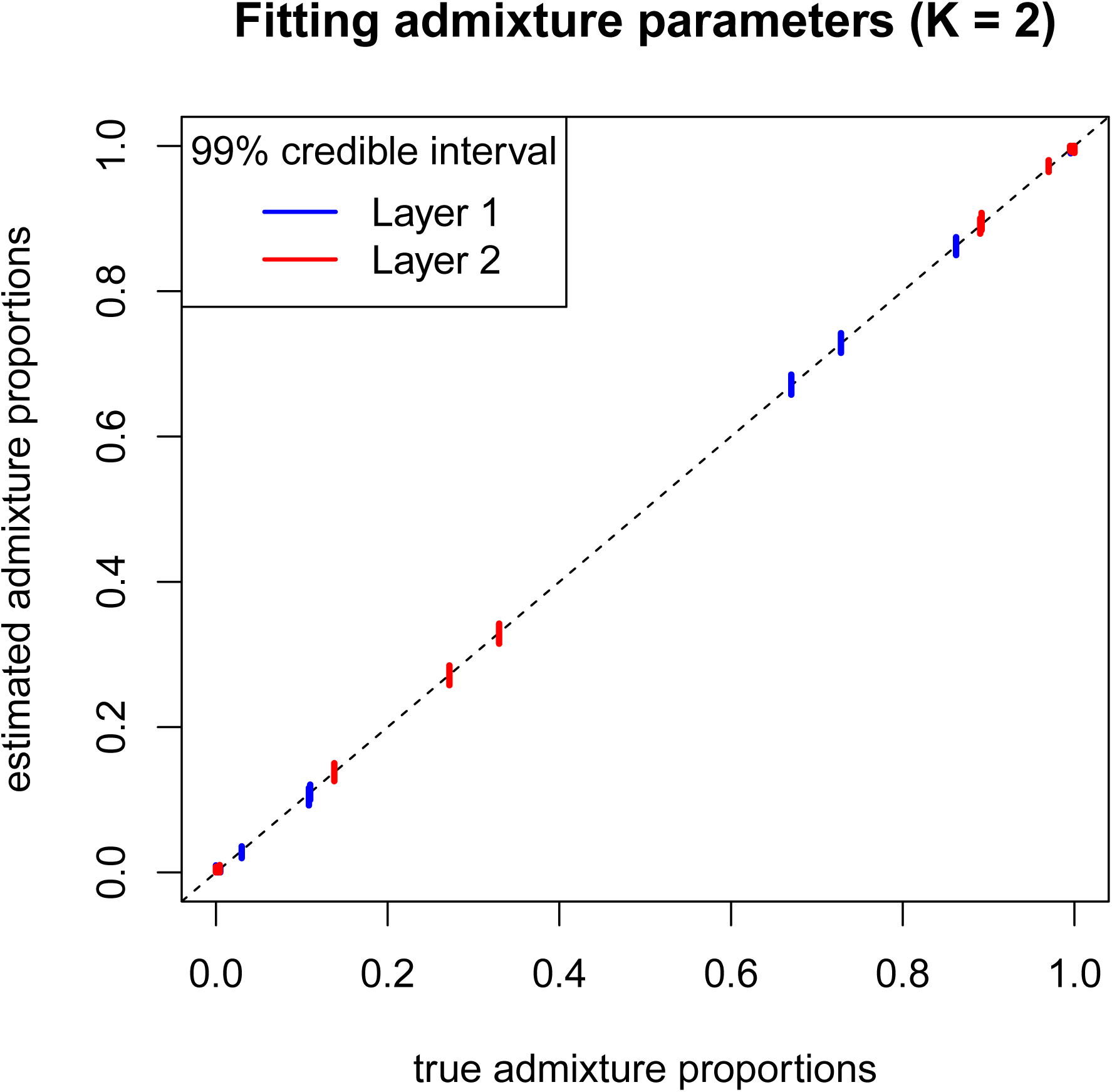
Plot of `conStruct` ability to correctly estimate admixture proportions on simulated data. Results are from an analysis with a spatial model using *K* = 2.

**Fig S9.**
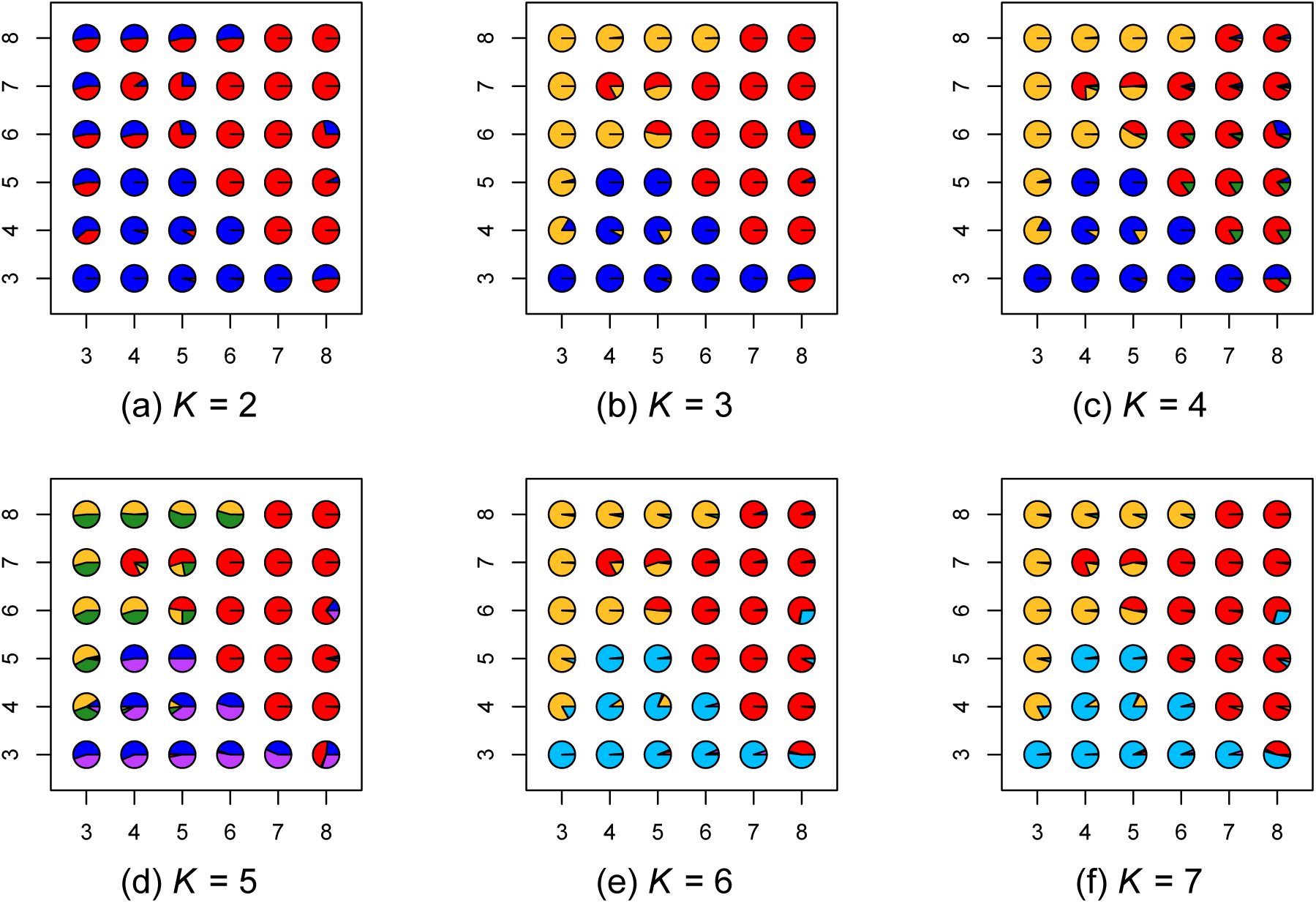
Map of admixture proportions estimated using a nonspatial model for *K* = 2 through 7. The data were simulated using three layers with nearest-neighbor symmetric migration between demes.

**Fig S10.**
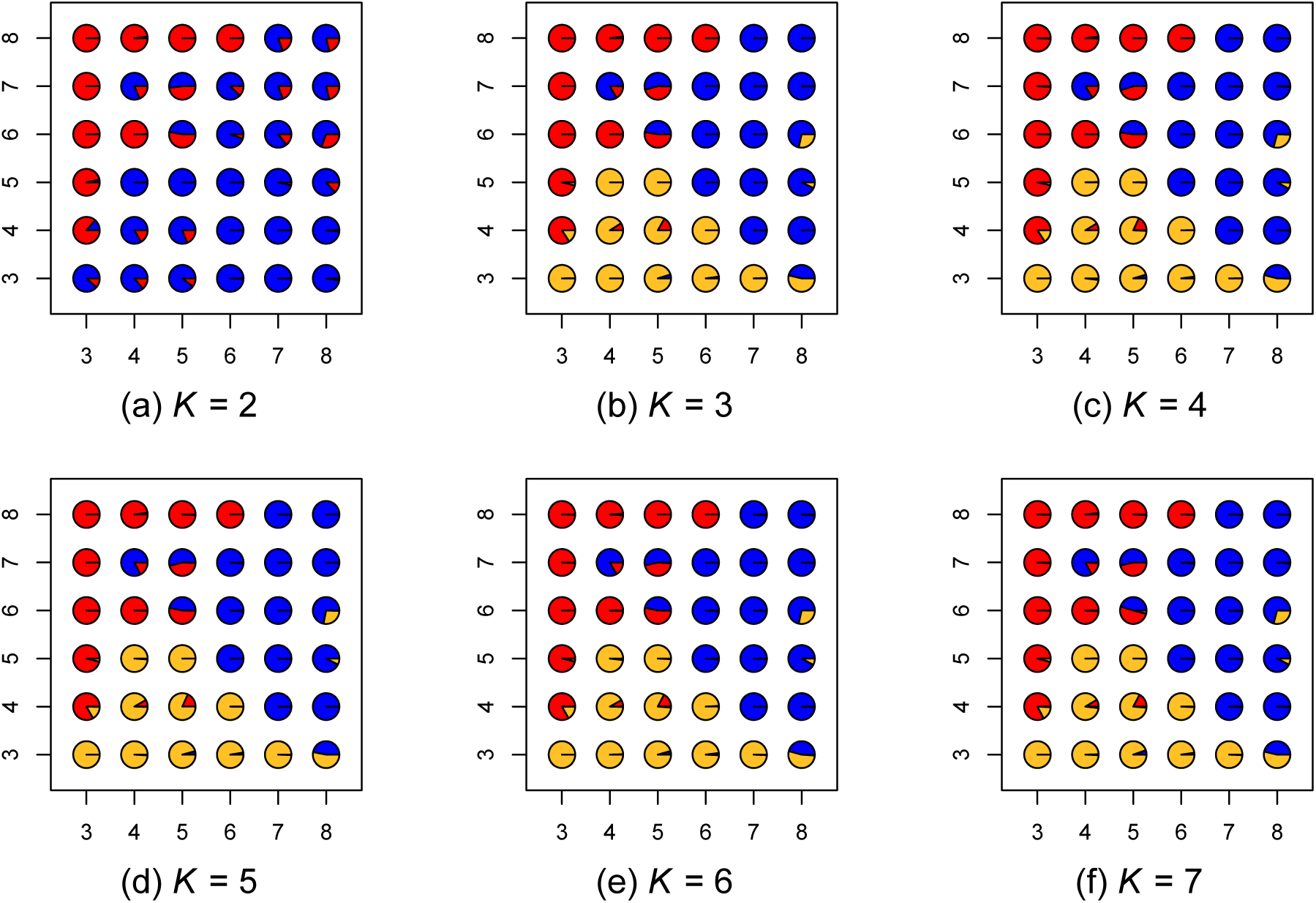
Map of admixture proportions estimated using a spatial model for *K* = 2 through 7. The data were simulated using three layers with nearest-neighbor symmetric migration between demes.

**Fig S11.**
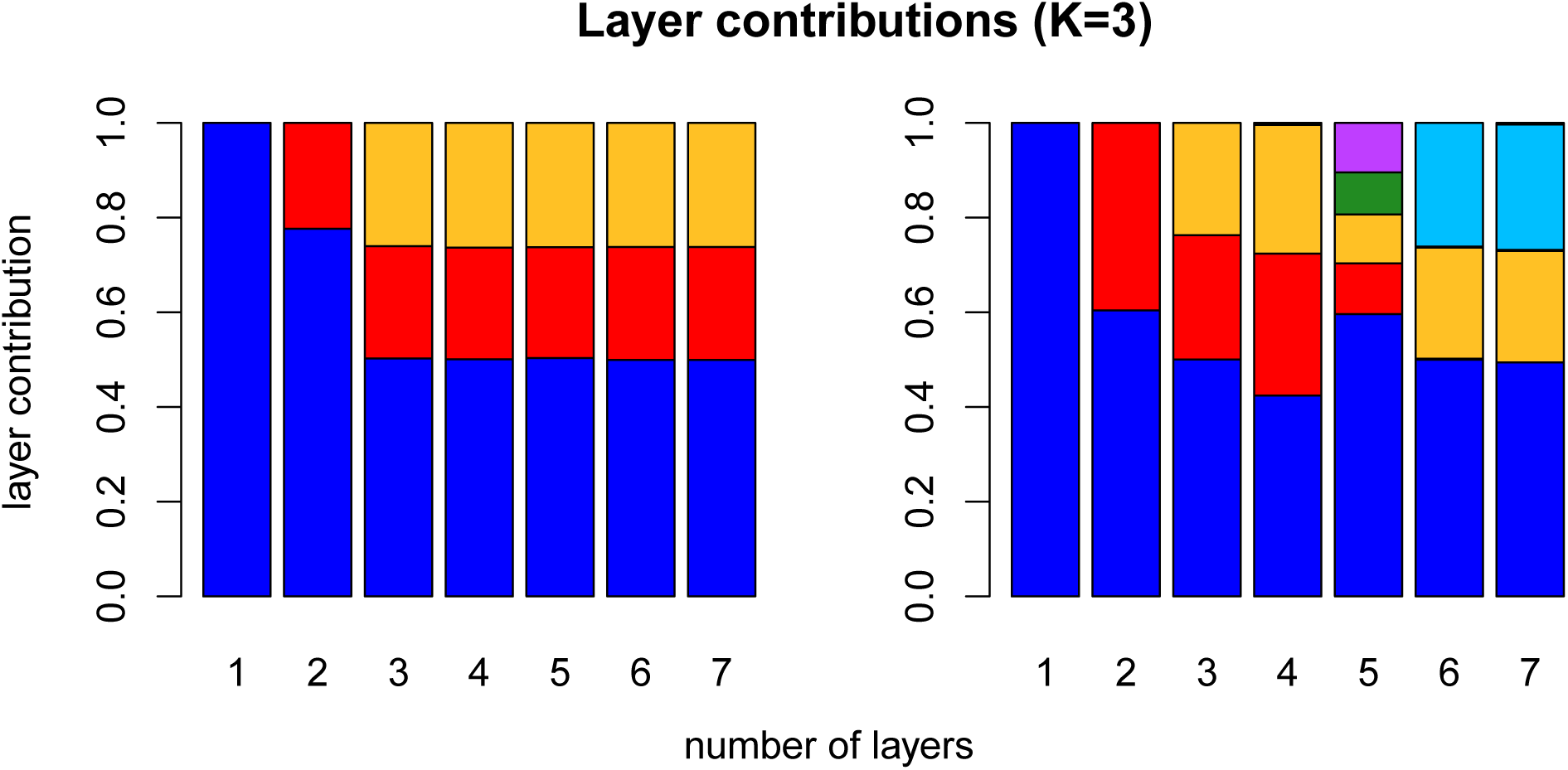
Layer/cluster contributions (i.e., how much total covariance is contributed by each layer/cluster), for all layers estimated in runs using *K* = 1 through 7 for the spatial model (left), and for all clusters using the nonspatial model (right). Data were simulated using *K* = 3. For each value of *K* along the x-axis, there are an equal number of contributions plotted.

**Fig S12.**
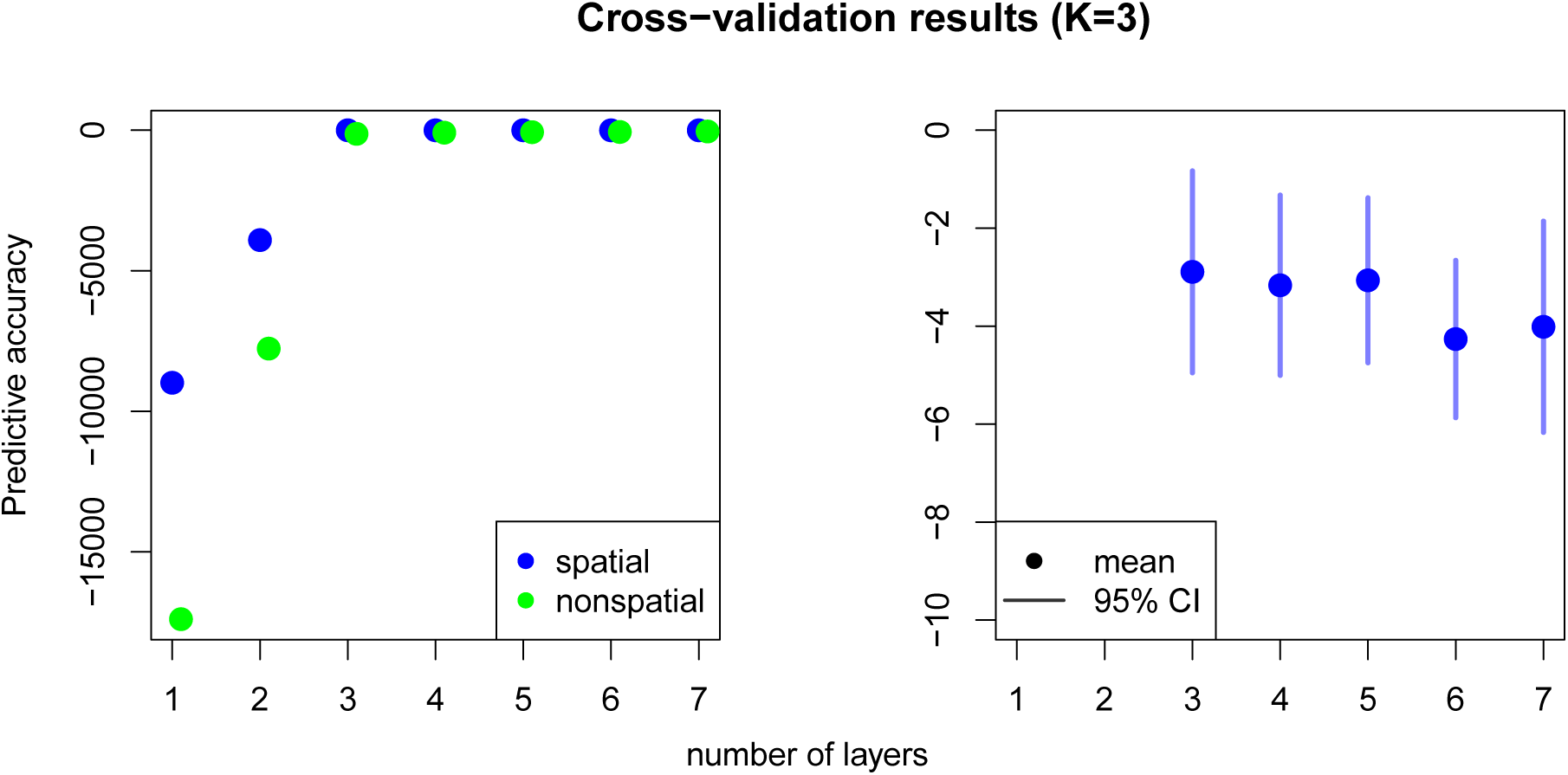
Cross-validation results for data simulated under *K* = 3, comparing the spatial and nonspatial `conStruct` models run with *K* = 1 through 7. The right panel zooms in on just the spatial cross-validation results.

**Fig S13.**
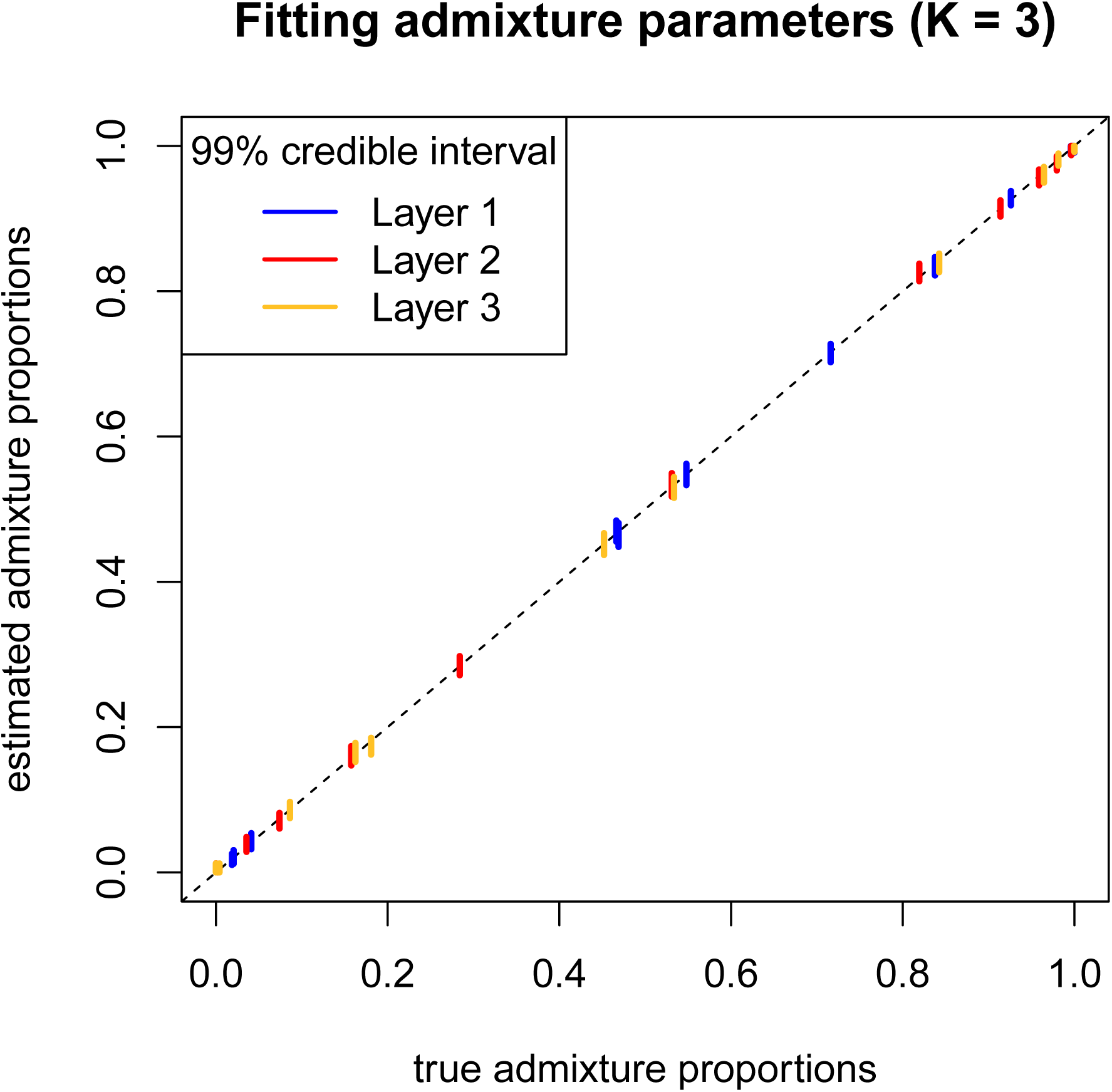
Plot of `conStruct` ability to correctly estimate admixture proportions on simulated data. Results are from an analysis with a spatial model using *K* = 3.

**Fig S14.**
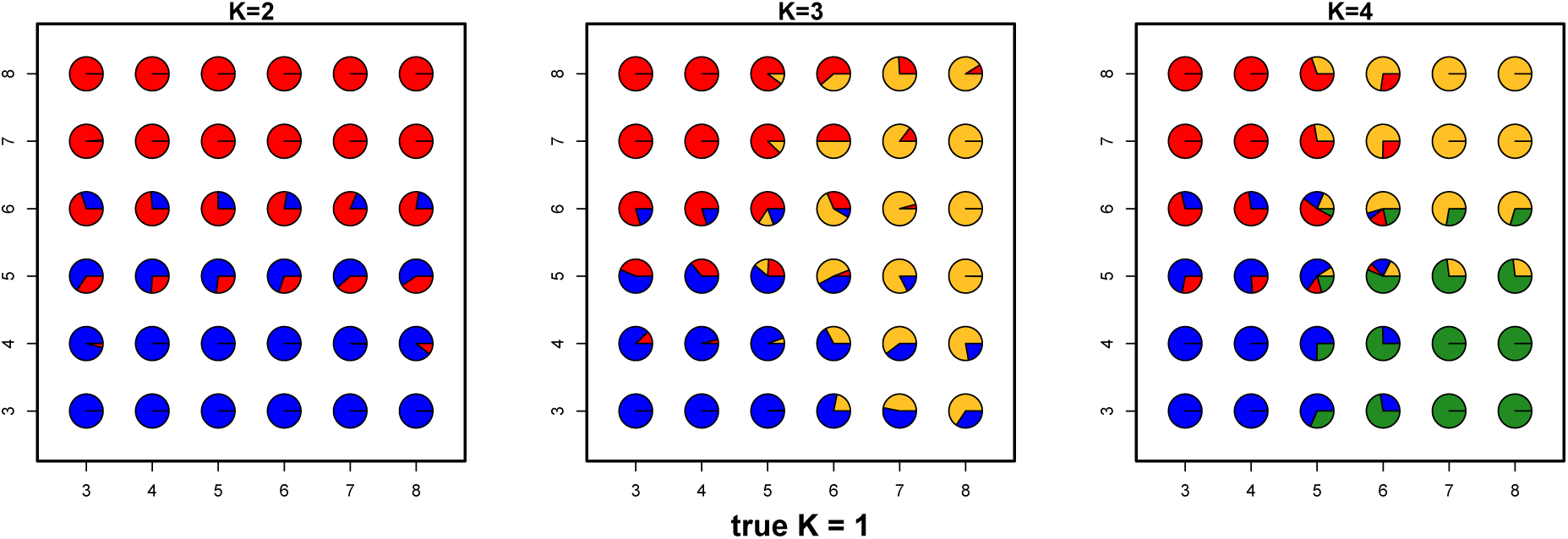
Map of admixture proportions estimated using fastSTRUCTURE [16] for *K* = 2 through 4. The data were simulated using one layer with nearest-neighbor symmetric migration between demes.

**Fig S15.**
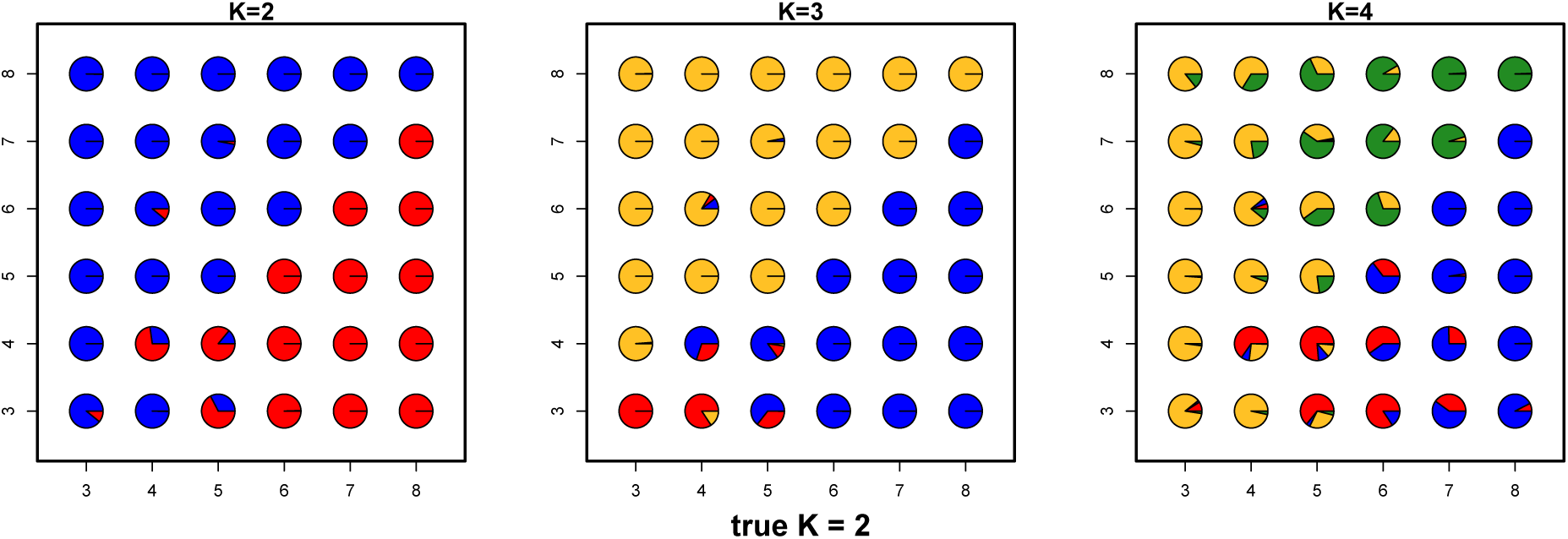
Map of admixture proportions estimated using fastSTRUCTURE [16] for *K* = 2 through 4. The data were simulated using two layers with nearest-neighbor symmetric migration between demes.

**Fig S16.**
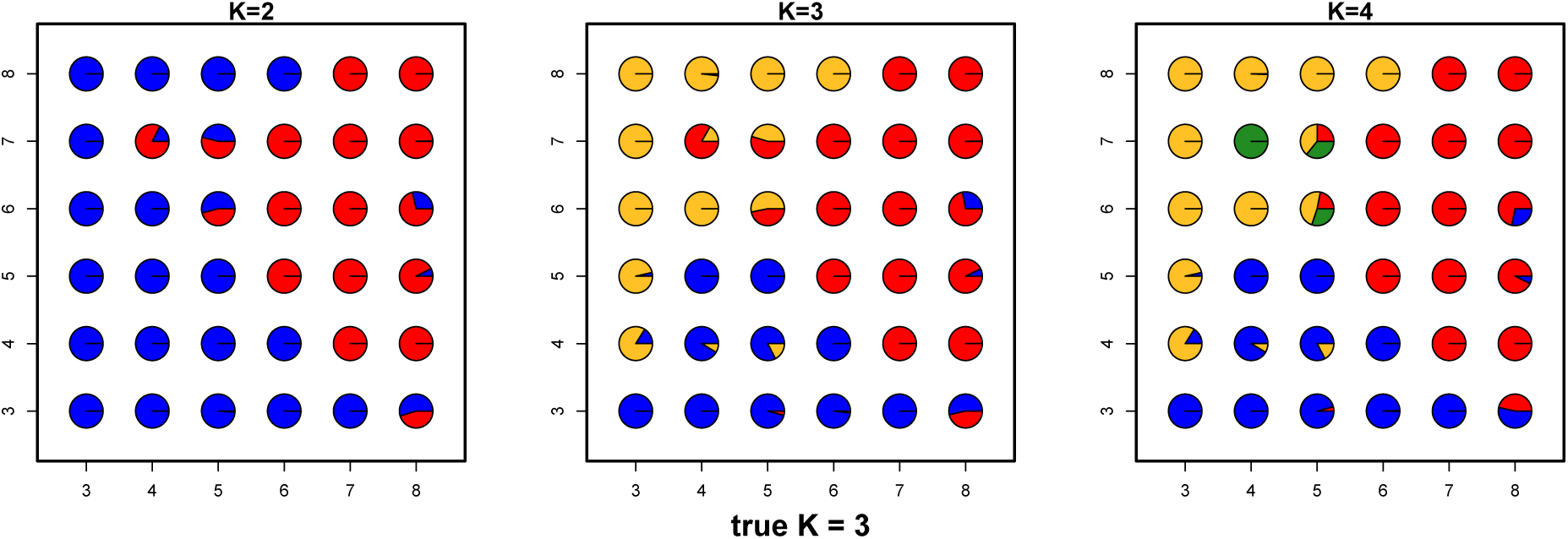
Map of admixture proportions estimated using fastSTRUCTURE [16] for *K* = 2 through 4. The data were simulated using three layers with nearest-neighbor symmetric migration between demes.

**Fig S17.**
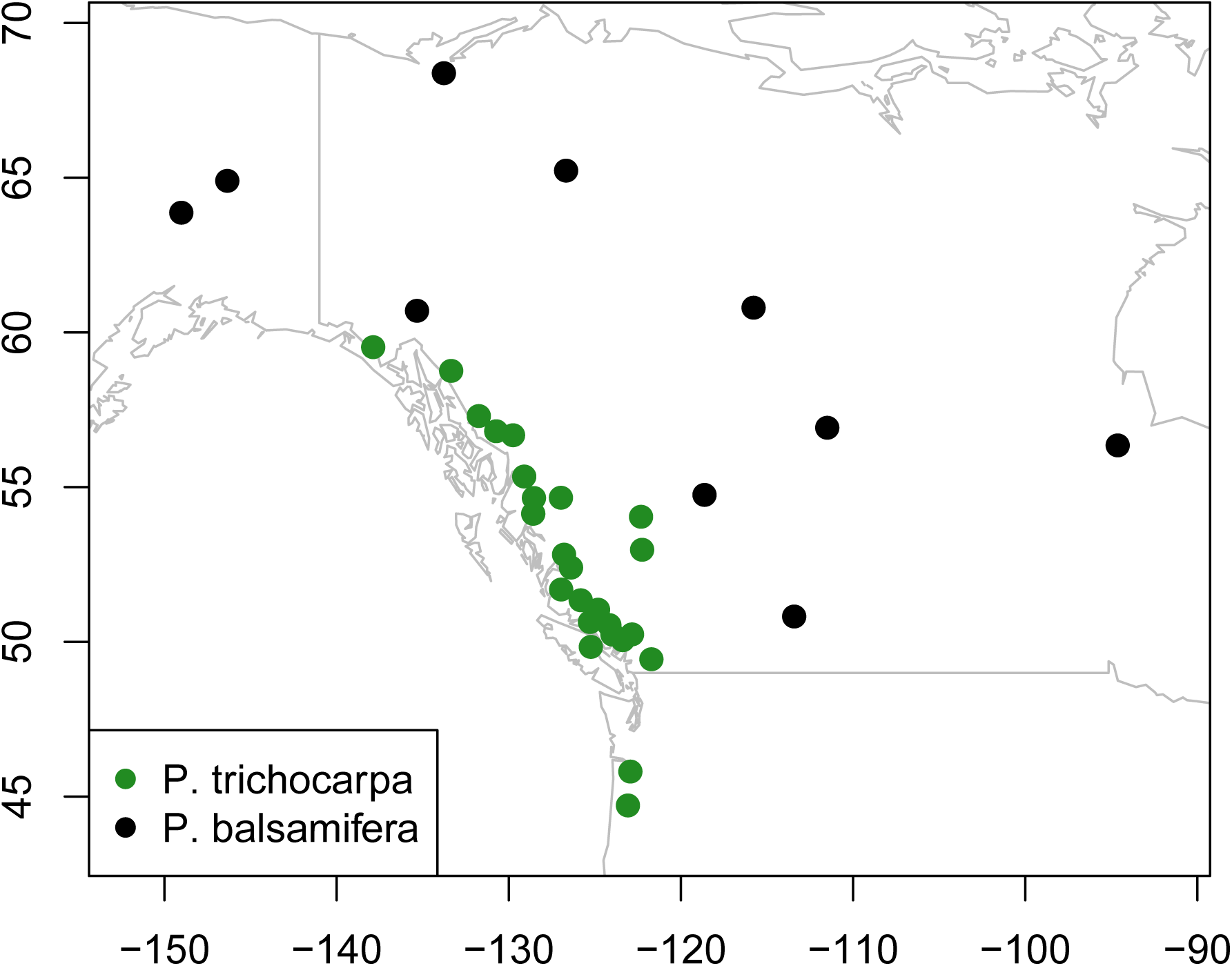
Map of the sampled *Populus* populations included in the analysis. The color of the sampling location denotes the putative species.

**Fig S18.**
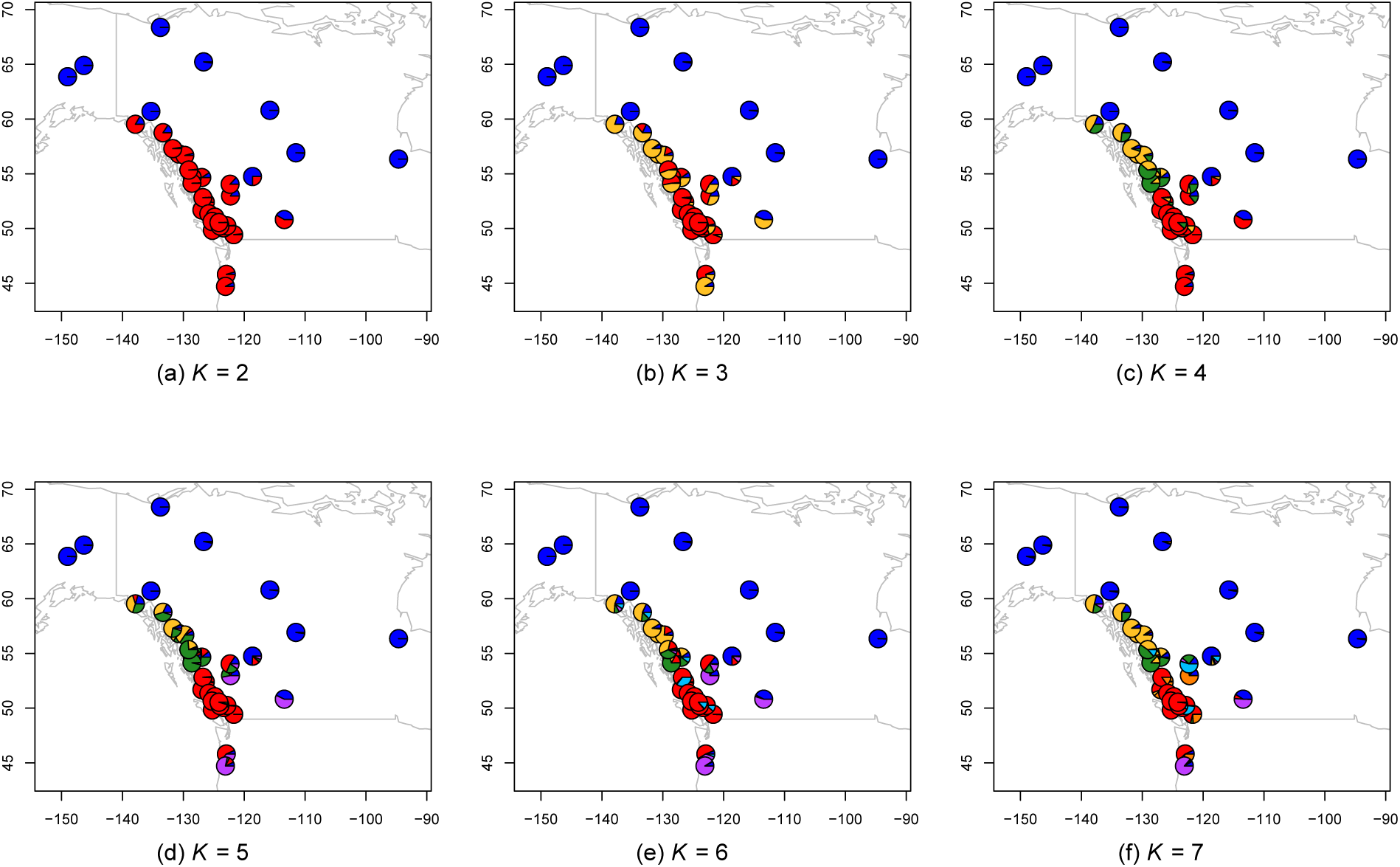
Maps of admixture proportions estimated for the *Populus* dataset using the spatial model for *K* = 2 through 7.

**Fig S19.**
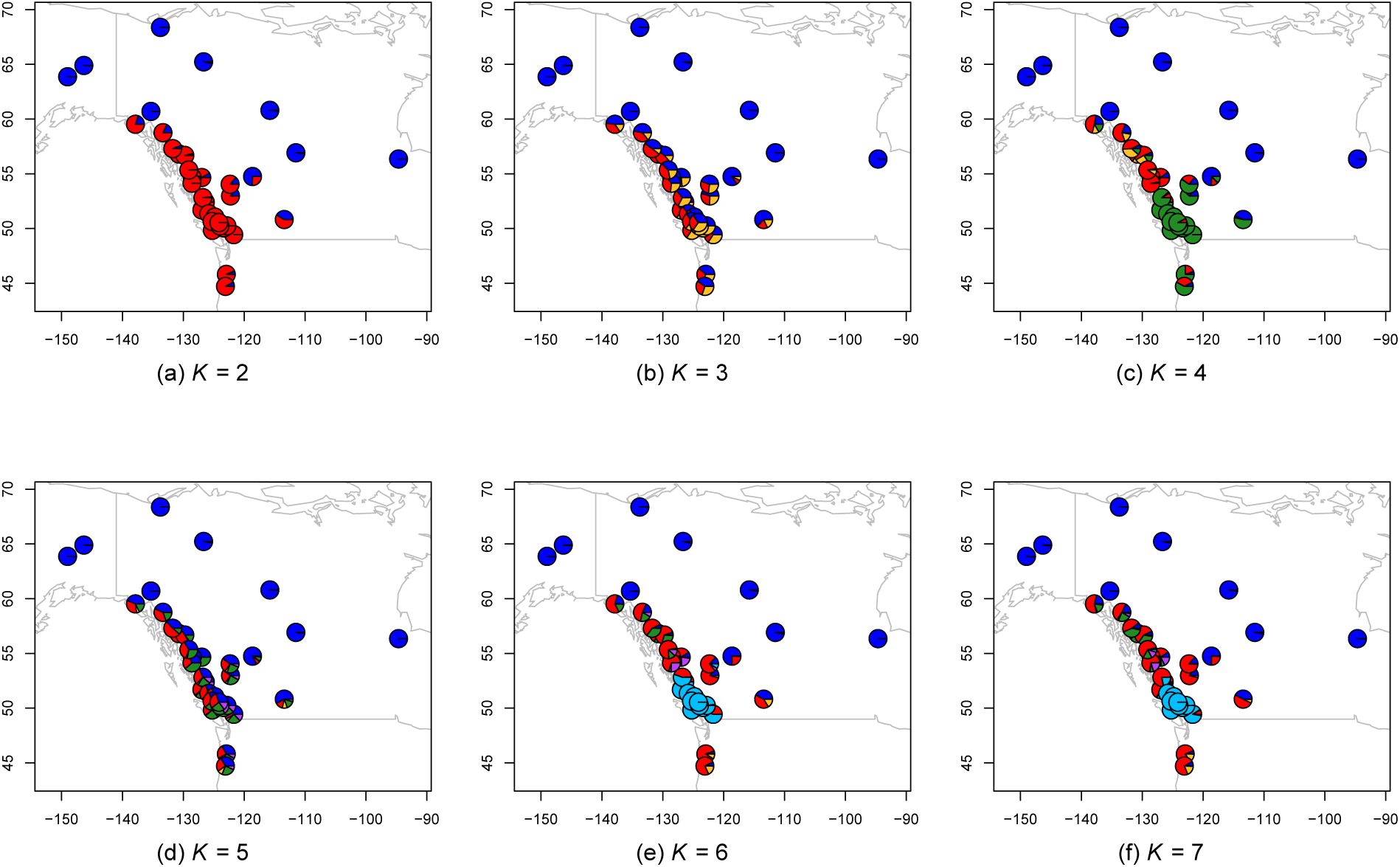
Maps of admixture proportions estimated for the *Populus* dataset using the nonspatial model for *K* = 2 through 7.

**Fig S20.**
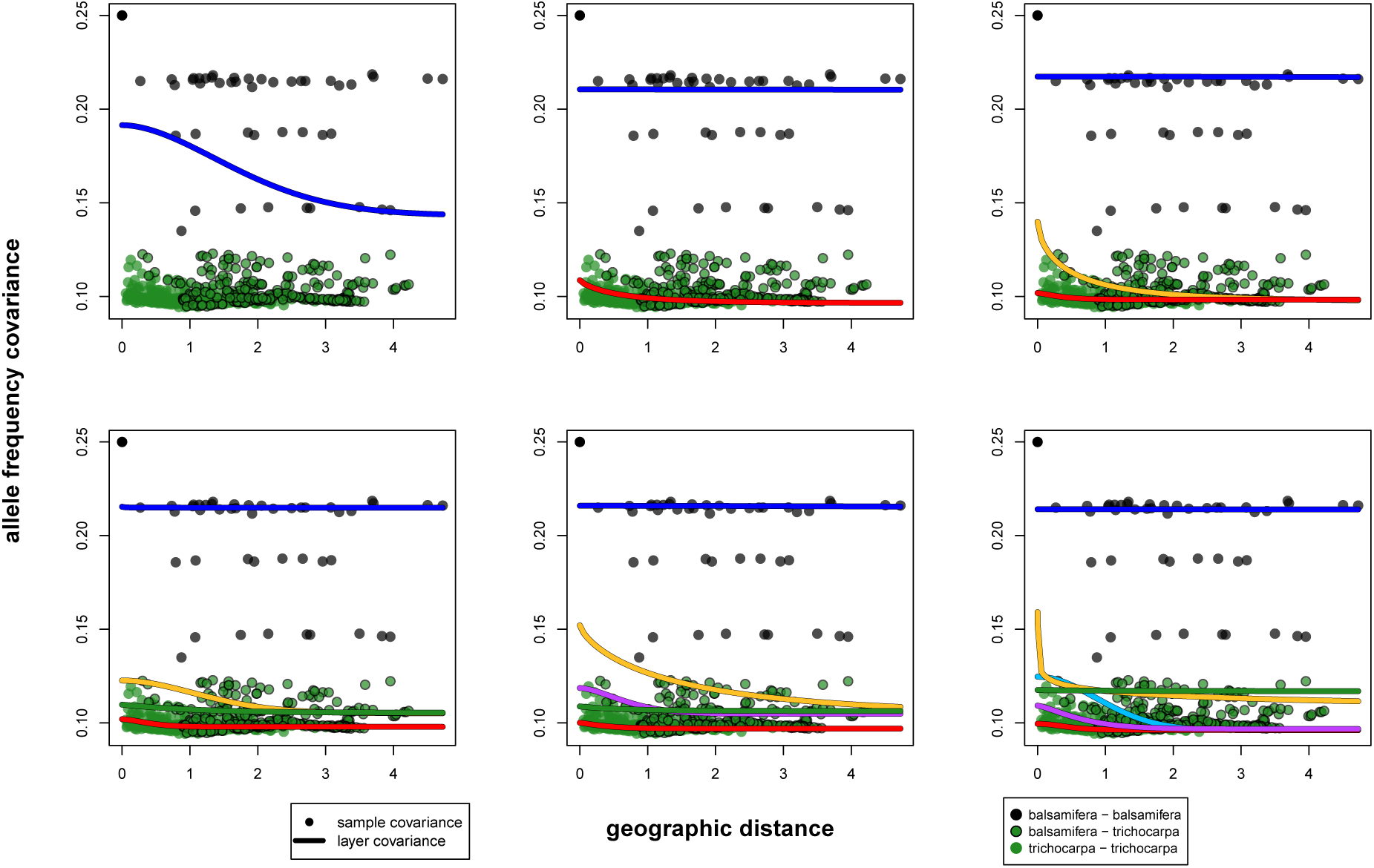
Plots showing the layer-specific parametric covariance curves estimated for the *Populus* data using the spatial `conStruct` model run with *K* = 1 through 6. Line colors are consistent with layer colors in Fig S18. Points are colored by the species they are a covariance between: black on black points are sample covariances between populations of *Populus balsamifera*; green on black points are sample covariances between *balsamifera* and *trichocarpa*; green on green points are sample covariances between *trichocarpa* and *trichocarpa*.

**Fig S21.**
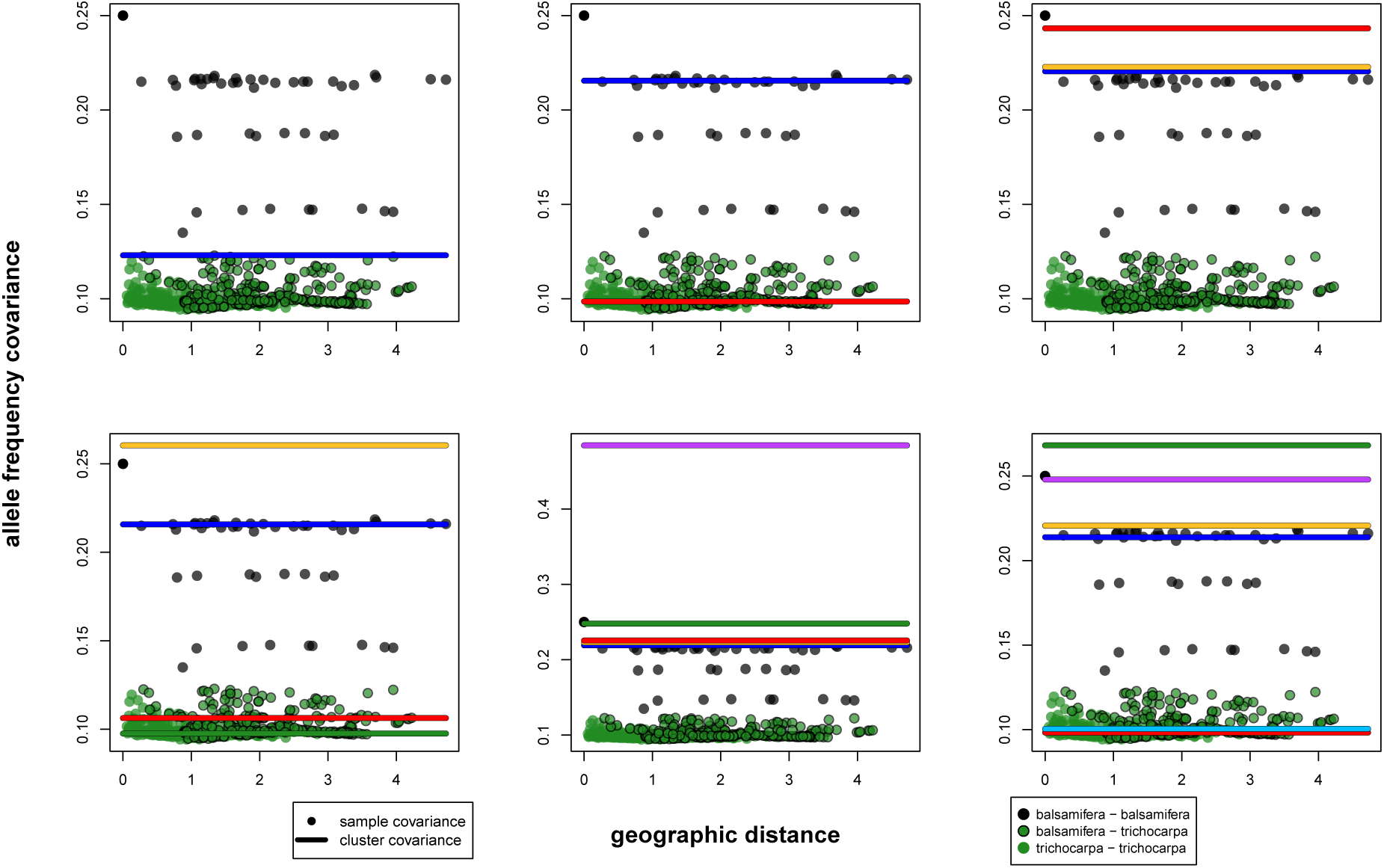
Plots showing the cluster-specific parametric covariances estimated for the *Populus* data using the nonspatial `conStruct` model run with *K* = 1 through 6. Line colors are consistent with cluster colors in Fig S19. Points are colored by the species they are a covariance between: black on black points are sample covariances between populations of *Populus balsamifera*; green on black points are sample covariances between *balsamifera* and *trichocarpa*; green on green points are sample covariances between *trichocarpa* and *trichocarpa*.

**Fig S22.**
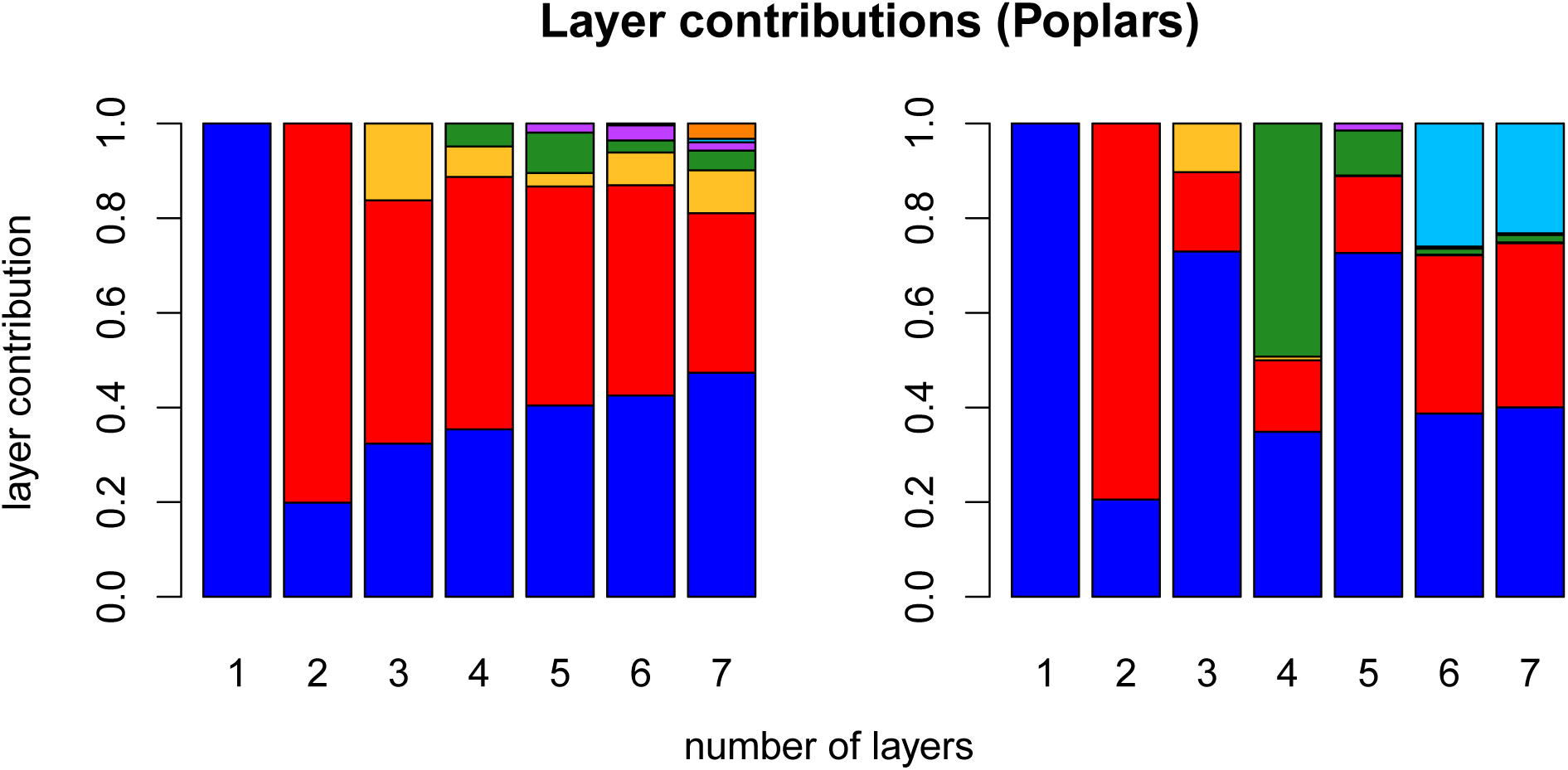
Layer/cluster contributions (i.e., how much total covariance is contributed by each layer/cluster), for all layers estimated in runs using *K* = 1 through 7 for the spatial model (left), and for all clusters using the nonspatial model (right). For each value of *K* along the x-axis, there are an equal number of contributions plotted. Colors are consistent with Figs S18, S20, S19, and S21.

**Fig S23.**
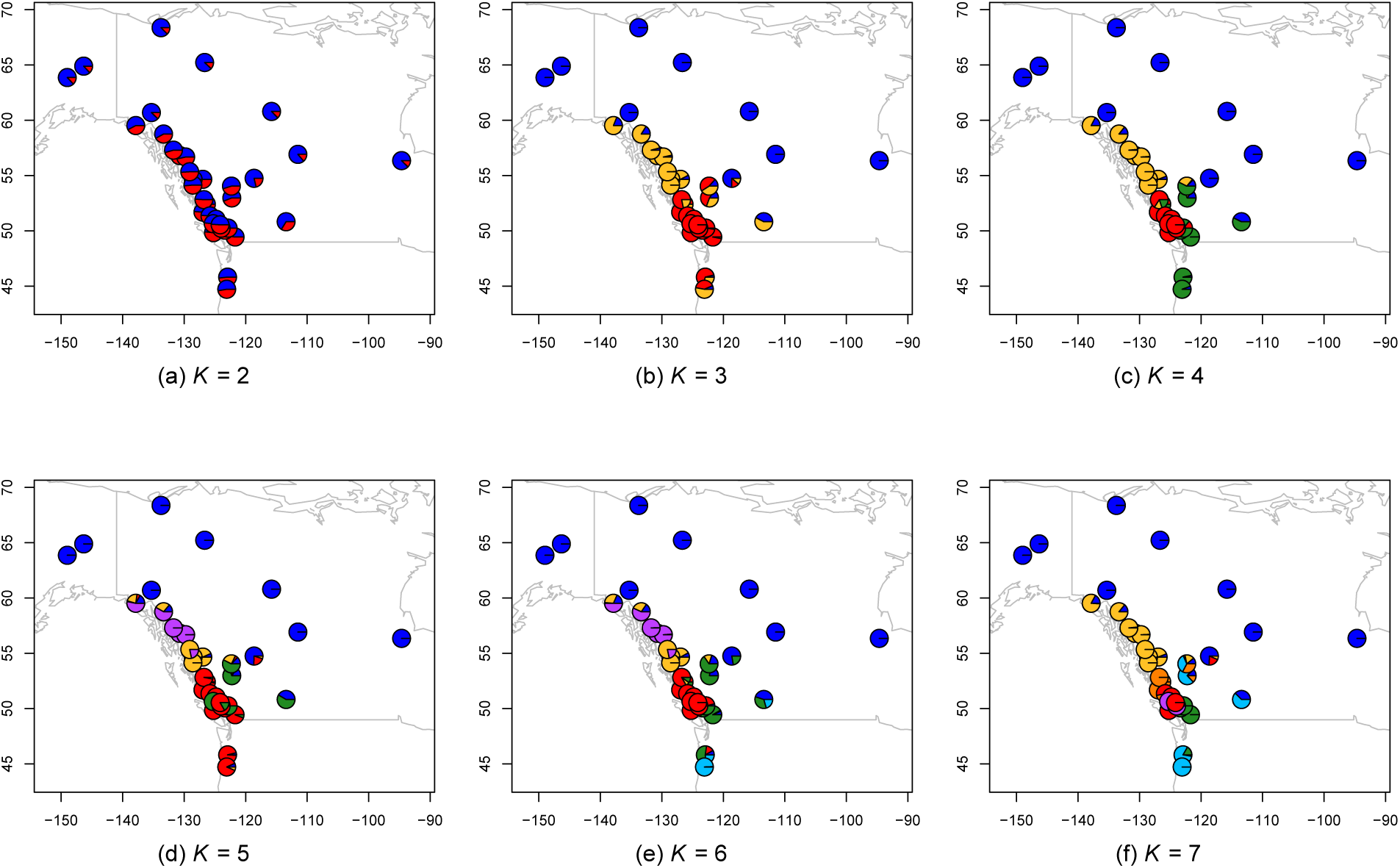
Maps of admixture proportions estimated for the *Populus* dataset using fastSTRUCTURE [16] for *K* = 2 through 7.

**Fig S24.**
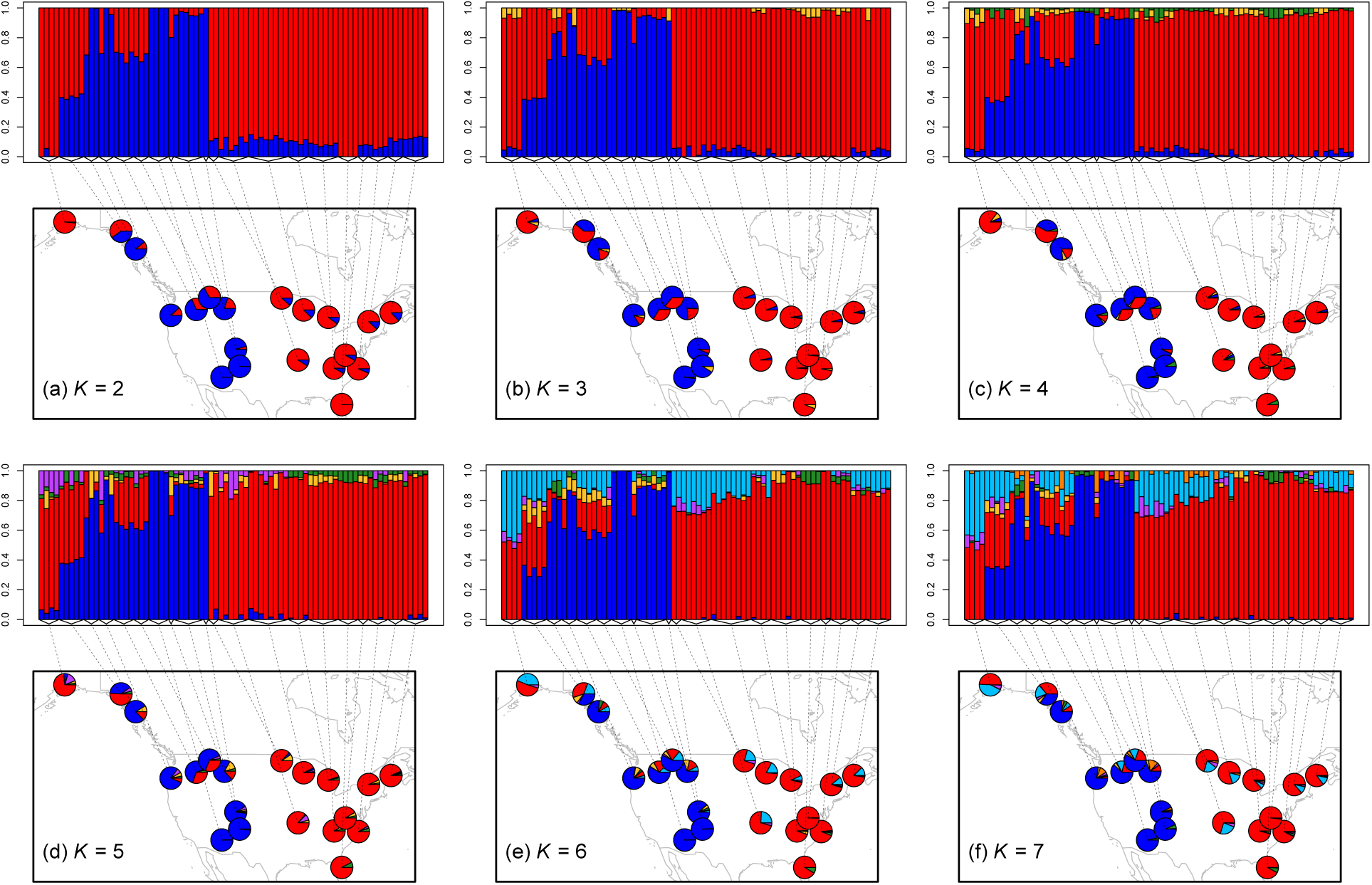
Map of admixture proportions estimated for the bear dataset using the spatial model for *K* = 2 through 7.

**Fig S25.**
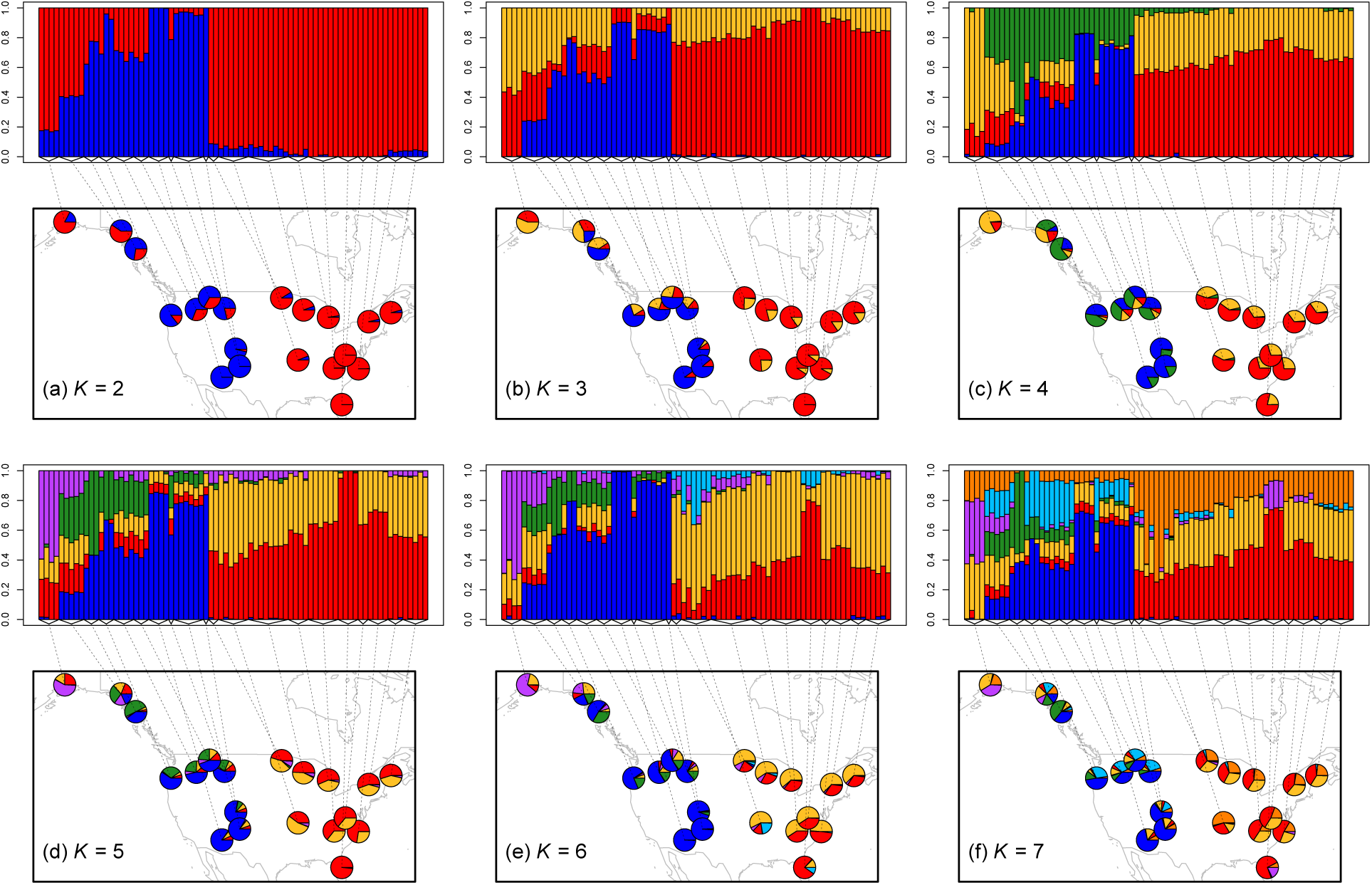
Map of admixture proportions estimated for the bear dataset using the nonspatial model for *K* = 2 through 7.

**Fig S26.**
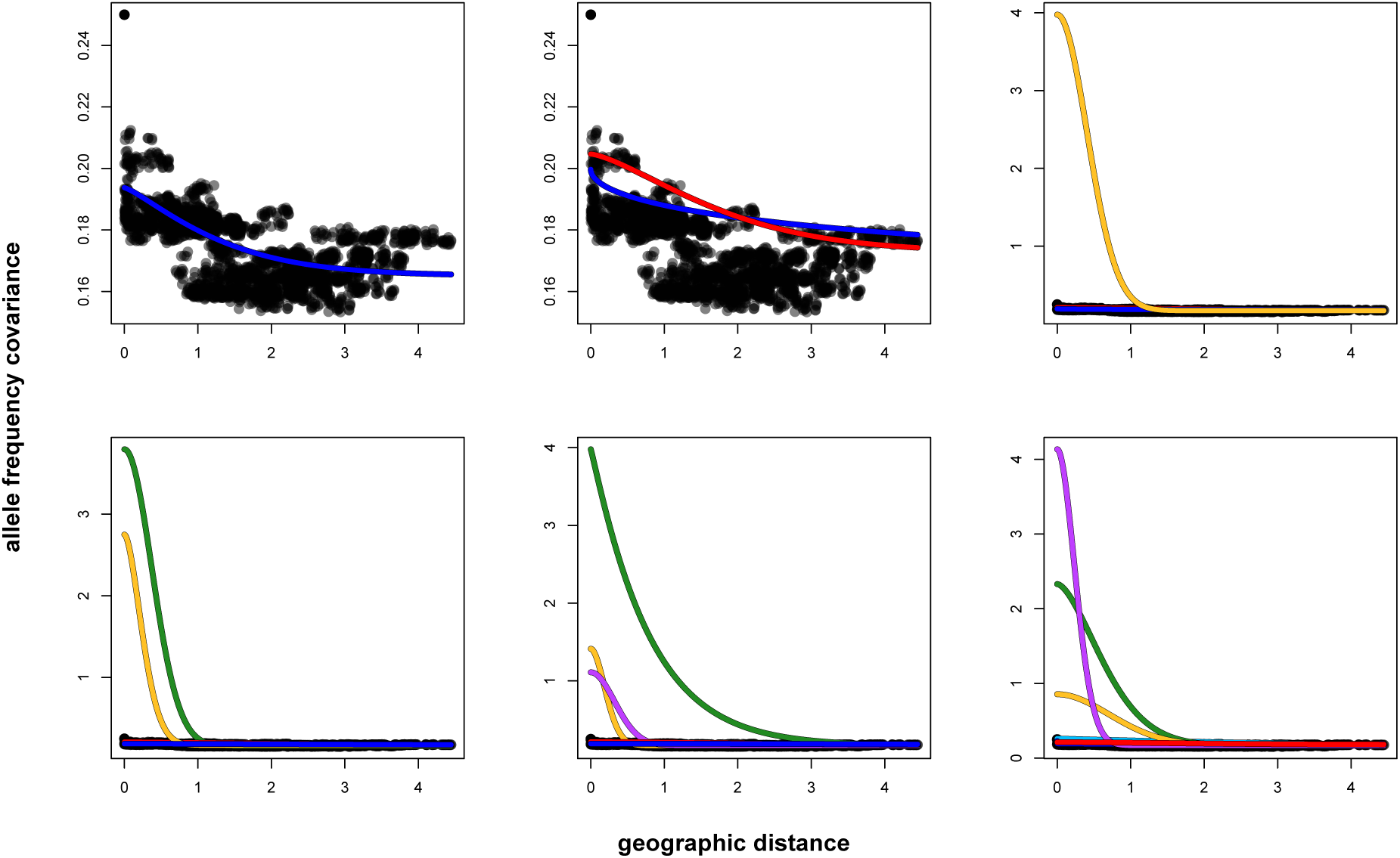
Plots showing the layer-specific parametric covariance curves estimated for the black bear data using the spatial `conStruct` model run with *K* = 1 through 6.

**Fig S27.**
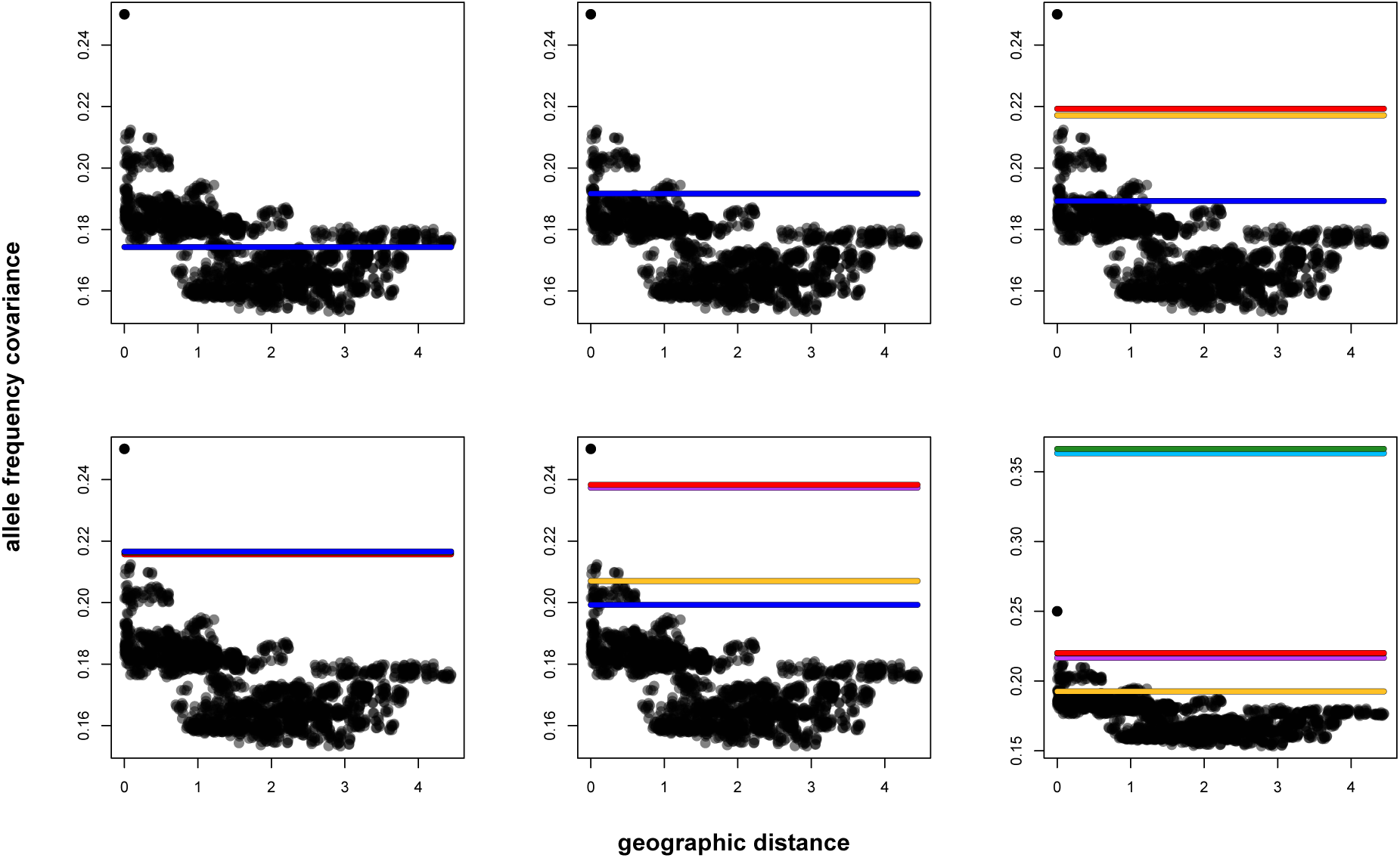
Plots showing the cluster-specific parametric covariances estimated for the black bear data using the nonspatial `conStruct` model run with *K* = 1 through 6.

**Fig S28.**
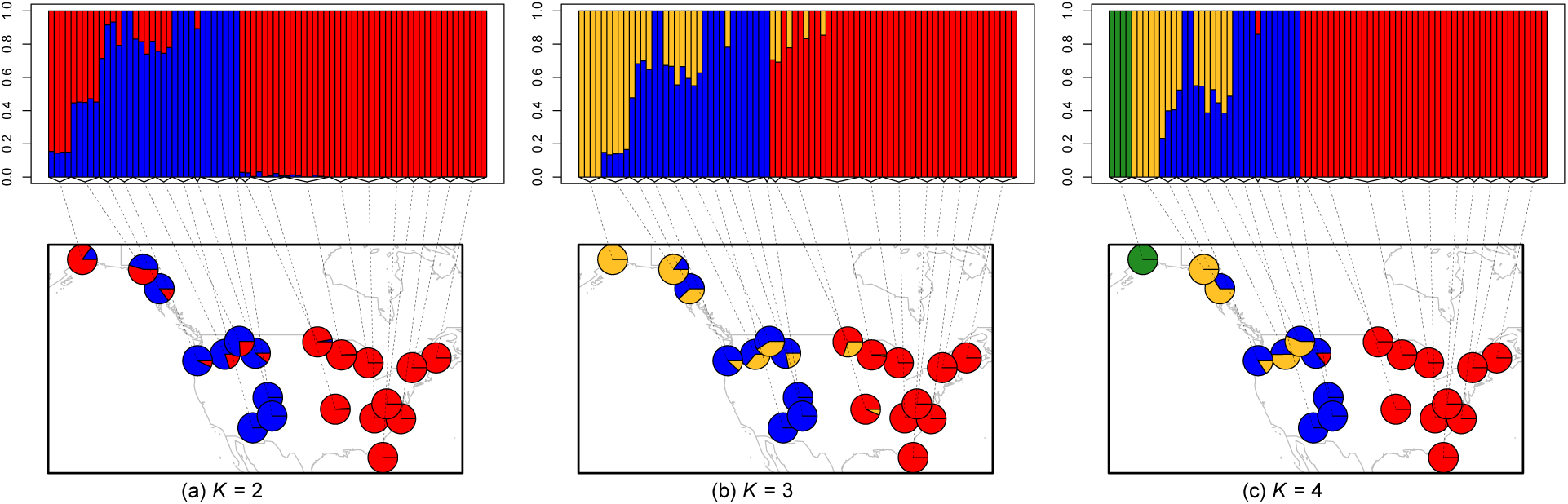
Maps of admixture proportions estimated for the black bear dataset using fastSTRUCTURE [16] for *K* = 2 through 4.

## References

1. Wright S. The genetical structure of populations. Annals of Eugenics. 1949;15(1):323–354.

2. Moritz C. Defining “Evolutionarily Significant Units” for conservation. Trends in Ecology and Evolution. 1994;9(10):373–375.

3. Waples R. Separating the wheat from the chaff: patterns of genetic differentiation in high gene flow species. Journal of Heredity. 1998;89(5):438–450.

4. Moritz C, Funk V, Sakai AK. Strategies to Protect Biological Diversity and the Evolutionary Processes That Sustain It. Systematic Biology. 2002;51(2):238–254.

5. Cavalli-Sforza LL, Piazza A. Analysis of evolution: Evolutionary rates, independence and treeness. Theoretical Population Biology. 1975;8(2):127–165.

6. Pickrell JK, Pritchard JK. Inference of population splits and mixtures from genome-wide allele frequency data. PLoS Genet. 2012;8(11):e1002967.

7. Meirmans P. GenoDive version 2.0 b14. Computer software distributed by the author Available from: http://www.bentleydrummernl/software/software/GenoDivehtml. 2009;.

8. Menozzi P, Piazza A, Cavalli-Sforza L. Synthetic maps of human gene frequencies in Europeans. Science. 1978;201(4358):786–792.

9. Novembre J, Stephens M. Interpreting principal component analyses of spatial population genetic variation. Nature genetics. 2008;40(5):646–649.

10. Price AL, Patterson NJ, Plenge RM, Weinblatt ME, Shadick NA, Reich D. Principal components analysis corrects for stratification in genome-wide association studies. Nat Genet. 2006;38(8):904–909.

11. Pritchard JK, Stephens M, Donnelly P. Inference of Population Structure Using Multilocus Genotype Data. Genetics. 2000;155(2):945–959.

12. Falush D, Stephens M, Pritchard JK. Inference of population structure using multilocus genotype data: linked loci and correlated allele frequencies. Genetics. 2003;164(4):1567–1587.

13. Hubisz MJ, Falush D, Stephens M, Pritchard JK. Inferring weak population structure with the assistance of sample group information. Molecular ecology resources. 2009;95:1322–32.

14. Alexander DH, Novembre J, Lange K. Fast model-based estimation of ancestry in unrelated individuals. Genome Research. 2009;19(9):1655–1664.

15. Lawson DJ, Hellenthal G, Myers S, Falush D. Inference of Population Structure using Dense Haplotype Data. PLoS Genet. 2012;8(1):e1002453.

16. Raj A, Stephens M, Pritchard JK. fastSTRUCTURE: Variational Inference of Population Structure in Large SNP Data Sets. Genetics. 2014;197(2):573–589.

17. Huelsenbeck JP, Andolfatto P. Inference of population structure under a Dirichlet process model. Genetics. 2007;175(4):1787–1802.

18. Corander J, Waldmann P, Sillanpää MJ. Bayesian Analysis of Genetic Differentiation Between Populations. Genetics. 2003;163(1):367–374.

19. Caye K, Jay F, Michel O, Francois O. Fast Inference of Individual Admixture Coefficients Using Geographic Data. bioRxiv. 2016;.

20. Guillot G, Mortier F, Estoup A. Geneland: a computer package for landscape genetics. Molecular Ecology Notes. 2005;5(3):712–715.

21. Wright S. Isolation by distance. Genetics. 1943;28(2):114–138.

22. Frantz AC, Cellina S, Krier A, Schley L, Burke T. Using spatial Bayesian methods to determine the genetic structure of a continuously distributed population: clusters or isolation by distance? Journal of Applied Ecology. 2009;46(2):493–505.

23. Kimura M, Weiss GH. The Stepping Stone Model of Population Structure and the Decrease of Genetic Correlation with Distance. Genetics. 1964;49(4):561–576.

24. Sawyer S. Results for the Stepping Stone Model for Migration in Population Genetics. The Annals of Probability. 1976;4(5):699–728.

25. Shiga T. Stepping stone models in population genetics and population dynamics. In: Stochastic processes in physics and engineering (Bielefeld, 1986). vol. 42 of Math. Appl. Dordrecht: Reidel; 1988. p. 345–355.

26. Malécot G. The Mathematics of Heredity. Freeman; 1969.

27. Slatkin M. Gene Flow in Natural Populations. Annual Review of Ecology and Systematics. 1985;16(1):393–430.

28. Epperson BK. Geographical Genetics. Monographs in Population Biology. Princeton University Press; 2003.

29. Barton NH, Depaulis F, Etheridge AM. Neutral Evolution in Spatially Continuous Populations. Theoretical Population Biology. 2002;61(1):31–48.

30. Barton NH, Etheridge AM, Véber A. Modelling evolution in a spatial continuum. Journal of Statistical Mechanics: Theory and Experiment. 2013;2013(01):P01002.

31. Meirmans PG. The trouble with isolation by distance. Molecular Ecology. 2012;21(12):2839–2846.

32. Sexton JP, Hangartner SB, Hoffmann AA. Genetic isolation by environment or distance: which pattern of gene flow is most common? Evolution. 2014;68(1):1–15.

33. Serre D, Pääbo S. Evidence for Gradients of Human Genetic Diversity Within and Among Continents. Genome Research. 2004;14(9):1679–1685.

34. Rosenberg NA, Mahajan S, Ramachandran S, Zhao C, Pritchard JK, Feldman MW. Clines, clusters, and the effect of study design on the inference of human population structure. PLoS Genet. 2005;1(6).

35. Wasser SK, Shedlock AM, Comstock K, Ostrander E, Mutayoba B, Stephens M. Assigning African elephant DNA to geographic region of origin: Applications to the ivory trade. PNAS. 2004;101(41):14847–52.

36. Bradburd GS, Ralph PL, Coop GM. Disentangling the effects of geographic and ecological isolation on genetic differentiation. Evolution. 2013;67(11):3258–3273.

37. Bradburd GS, Ralph PL, Coop GM. A Spatial Framework for Understanding Population Structure and Admixture. PLoS Genet. 2016;12(1):1–38.

38. McVean G. A Genealogical Interpretation of Principal Components Analysis. PLoS Genet. 2009;5(10):e1000686.

39. Petkova D, Novembre J, Stephens M. Visualizing spatial population structure with estimated effective migration surfaces. Nature genetics. 2016;48(1):94–100C.

40. Linck EB, Battey CJ. Minor allele frequency thresholds strongly affect population structure inference with genomic datasets. bioRxiv. 2017;.

41. Diggle PJ, Tawn JA, Moyeed RA. Model-based geostatistics. Jounal of the Royal Statistical Society Series C (Applied Statistics). 1998;47(3):299–350.

42. Patterson N, Moorjani P, Luo Y, Mallick S, Rohland N, Zhan Y, et al. Ancient Admixture in Human History. Genetics. 2012;192(3):1065–1093.

43. Peter BM. Admixture, Population Structure and F-Statistics. Genetics. 2016;.

44. Carpenter B. Stan: A Probabilistic Programming Language. Journal of Statistical Software. 2015;.

45. Hoffman MD, Gelman A. The No-U-Turn Sampler: Adaptively Setting Path Lengths in Hamiltonian Monte Carlo. Journal of Machine Learning Research. 2014;.

46. Stan Development Team. Stan: A C++ Library for Probability and Sampling, Version 2.10.0; 2015.

47. Stan Development Team. RStan: the R interface to Stan, Version 2.10.1; 2016.

48. Engelhardt BE, Stephens M. Analysis of Population Structure: A Unifying Framework and Novel Methods Based on Sparse Factor Analysis. PLOS Genetics. 2010;6(9):1–12.

49. Evanno G, Regnaut S, Goudet J. Detecting the number of clusters of individuals using the software structure: a simulation study. Molecular Ecology. 2005;14(8):2611–2620.

50. Verity R, Nichols R. Estimating K in Genetic Mixture Models. bioRxiv. 2015;.

51. Alexander DH, Lange K. Enhancements to the ADMIXTURE Algorithm for Individual Ancestry Estimation. BMC Bioinformatics. 2011;12:246.

52. Hudson RR. Generating samples under a Wright-Fisher neutral model of genetic variation. Bioinformatics. 2002;18(2):337–338.

53. Eckenwalder JE. Natural intersectional hybridization between North American species of Populus (Salicaceae) in sections Aigeiros and Tacamahaca. II. Taxonomy. Canadian Journal of Botany. 1984;62(2):325–335.

54. Cronk QCB. Plant eco-devo: the potential of poplar as a model organism. New Phytologist. 2005;166(1):39–48.

55. Geraldes A, Farzaneh N, Grassa CJ, McKown AD, Guy RD, Mansfield SD, et al. Landscape genomics of Populus trichocarpa: the role of hybridization, limited gene flow, and natural selection in shaping patterns of population structure. Evolution. 2014;68(11):3260–3280.

56. Suarez-Gonzalez A, Hefer CA, Christe C, Corea O, Lexer C, Cronk QCB, et al. Genomic and functional approaches reveal a case of adaptive introgression from Populus balsamifera (balsam poplar) in P. trichocarpa (black cottonwood). Molecular Ecology. 2016;25(11):2427–2442.

57. Slavov GT, DiFazio SP, Martin J, Schackwitz W, Muchero W, Rodgers-Melnick E, et al. Genome resequencing reveals multiscale geographic structure and extensive linkage disequilibrium in the forest tree Populus trichocarpa. New Phytologist. 2012;196(3):713–725.

58. McKown AD, Guy RD, Klapste J, Geraldes A, Friedmann M, Cronk QCB, et al. Geographical and environmental gradients shape phenotypic trait variation and genetic structure in Populus trichocarpa. New Phytologist. 2014;201(4):1263–1276.

59. Keller SR, Olson MS, Silim S, Schroeder W, Tiffin P. Genomic diversity, population structure, and migration following rapid range expansion in the Balsam Poplar, Populus balsamifera. Molecular Ecology. 2010;19(6):1212–1226.

60. Xie CY, Ying CC, Yanchuk AD, Holowachuk DL. Ecotypic mode of regional differentiation caused by restricted gene migration: a case in black cottonwood (Populus trichocarpa) along the Pacific Northwest coast. Canadian Journal of Forest Research. 2009;39(3):519–525.

61. Chang-Yi X, R CM, C YC. Ecotypic mode of regional differentiation of black cottonwood (Populus trichocarpa) due to restricted gene migration: further evidence from a field test on the northern coast of British Columbia. Canadian Journal of Forest Research. 2012;42(2):400–405.

62. Zhou L, Holliday JA. Targeted enrichment of the black cottonwood (Populus trichocarpa) gene space using sequence capture. BMC Genomics. 2012;13(1):703.

63. Wooding S, Ward R. Phylogeography and pleistocene evolution in the North American black bear. Molecular Biology and Evolution. 1997;14(11):1096–1105.

64. Byun SA, Koop BF, Reimchen TE. North American Black Bear mtDNA Phylogeography: Implications for Morphology and the Haida Gwaii Glacial Refugium Controversy. Evolution. 1997;51(5):1647–1653.

65. Stone KD, Cook JA. Phylogeography of black bears (Ursus americanus) of the Pacific Northwest. Canadian Journal of Zoology. 2000;78(7):1218–1223.

66. Puckett EE, Etter PD, Johnson EA, Eggert LS. Phylogeographic Analyses of American Black Bears (Ursus americanus) Suggest Four Glacial Refugia and Complex Patterns of Postglacial Admixture. Molecular Biology and Evolution. 2015;32(9):2338–2350.

67. Woodbury MA. Inverting modified matrices. Statistical Research Group, Memo. Rep. no. 42. Princeton University, Princeton, N. J.; 1950.

68. Sherman J, Morrison WJ. Adjustment of an Inverse Matrix Corresponding to a Change in One Element of a Given Matrix. Ann Math Statist. 1950;21(1):124–127.

69. Patterson N, Price AL, Reich D. Population Structure and Eigenanalysis. PLoS Genet. 2006;2(12):e190.

70. Falush D, van Dorp L, Lawson D. A tutorial on how (not) to over-interpret STRUCTURE/ADMIXTURE bar plots. bioRxiv. 2016;.

71. Irwin DE, Bensch S, Price TD. Speciation in a ring. Nature. 2001;409(6818):333–337.

72. Wake DB, Schneider CJ. Taxonomy of the Plethodontid Salamander Genus Ensatina. Herpetologica. 1998;54(2):pp. 279–298.

73. Lazaridis I, Patterson N, Mittnik A, Renaud G, Mallick S, Kirsanow K, et al. Ancient human genomes suggest three ancestral populations for present-day Europeans. Nature. 2014;513(7518):409–413.

74. Haak W, Lazaridis I, Patterson N, Rohland N, Mallick S, Llamas B, et al. Massive migration from the steppe was a source for Indo-European languages in Europe. Nature. 2015;522(7555):207–211.

75. Slatkin M, Racimo F. Ancient DNA and human history. Proceedings of the National Academy of Sciencess. 2016;113(23):6380–6387.

76. Nielsen R, Akey JM, Jakobsson M, Pritchard JK, Tishkoff S, Willerslev E. Tracing the peopling of the world through genomics. Nature. 2017;541(7637):302–310.

77. Schraiber J. Assessing the relationship of ancient and modern populations. bioRxiv. 2017;.

78. Picard RR, Cook RD. Cross-Validation of Regression Models. Journal of the American Statistical Association. 1984;79(387):575–583.

